# One-shot tagging during wake and cueing during sleep with spatiotemporal patterns of transcranial electrical stimulation can boost long-term metamemory of individual episodes in humans

**DOI:** 10.1101/672378

**Authors:** Praveen K. Pilly, Steven W. Skorheim, Ryan J. Hubbard, Nicholas A. Ketz, Shane M. Roach, Aaron P. Jones, Bradley Robert, Natalie B. Bryant, Itamar Lerner, Arno Hartholt, Teagan S. Mullins, Jaehoon Choe, Vincent P. Clark, Michael D. Howard

**Author notes:** These authors contributed equally to this work (Steven W. Skorheim, Ryan J. Hubbard).

## Abstract

Targeted memory reactivation (TMR) during slow-wave oscillations (SWOs) in non-rapid eye movement (NREM) sleep has been demonstrated with sensory cues to achieve about 5-12% improvement in post-nap memory performance on simple laboratory tasks. But prior work has neither addressed the one-shot aspect of episodic memory acquisition, nor dealt with the presence of interference from ambient environmental cues in real-world settings for the sensory cues. Moreover, TMR with sensory cues may not be scalable to the multitude of experiences over one’s lifetime. We designed a novel non-invasive paradigm that tags one-shot experiences of minute-long naturalistic episodes within immersive virtual reality (VR) with unique spatiotemporal amplitude-modulated patterns (STAMPs) of transcranial electrical stimulation (tES) and cues them during SWOs. In particular, we demonstrate that these STAMPs can be re-applied as brief pulses to temporally coincide with UP states of SWOs (0.4167 – 1 s) on two consecutive nights to achieve about 20% improvement in the metamemory of targeted episodes at 48 hours after the one-shot viewing, compared to the control episodes. Post-sleep metamemory of the targeted episodes was driven by an interaction between their pre-sleep metamemory and the number of STAMP applications for those episodes during sleep. Overnight metamemory improvements were mediated by spectral power increases from 6.18 to 6.7 s following the offset of STAMPs in the slow-spindle band (9-12 Hz) for left temporal areas in the scalp electroencephalography (EEG) during sleep. These results prescribe an optimal strategy to leverage STAMPs for boosting metamemory and suggest that real-world episodic memories can be modulated in a targeted manner even with coarser, non-invasive spatiotemporal stimulation.

## Introduction

Applied neuroscience aims to develop technologies to affect behavior by modulating brain activity at the right spatiotemporal scale. The ability to recall previously experienced events and to introspect about them are important aspects of our daily living. However, a mechanistic understanding of how episodic memories of real-world events are formed, recalled, and monitored in the human brain is still lacking. This is especially challenging given that such events are typically experienced only once. Metamemory is an executive function that monitors and judges the ability to recall memories accurately (Nelson & Narens, 1990), such as when providing eyewitness testimony in a criminal case and determining which learning material needs more study. One is said to have higher metamemory when their recall accuracy is proportional to their subjective confidence (i.e., more confident when correct and less confident when wrong). In other words, metamemory measures the ability to introspect and discriminate between correct and incorrect memory recalls, avoiding either over- or under-confidence (Galvin et al., 2003; Fleming & Lau, 2014). The neural mechanisms underlying memory monitoring and control are known to involve dorsolateral prefrontal, parietal, and cingulate cortices (Chua et al., 2009), and likely work in concert with those involved in the encoding, consolidation, and recall of the memory content (Nelson & Narens, 1990).

Hippocampus plays an important role in the online rapid encoding of episodic memories for short-term storage, which subsequently drives offline slow consolidation for long-term storage in distributed neocortical areas (McClelland et al., 1995; Buzsaki, 1996). It has been suggested that any intervention to enhance specific episodic memories needs to operate with high spatiotemporal resolution in the hippocampus (Hampson et al., 2018; Suthana & Fried, 2014). But it is also possible for widespread neocortical activations to trigger coordinated memory replays in the hippocampus (Ji & Wilson, 2007; Rothschild et al., 2017). Consistent with this latter view, there have been a number of demonstrations of targeted memory reactivation (TMR) in animals and humans using olfactory and auditory cues to modulate the ability to learn contexts and individual memories (e.g., Rasch et al., 2007; Rudoy et al., 2009; Antony et al., 2012; Bendor & Wilson, 2012). But these studies have neither addressed the one-shot aspect of episodic memory acquisition, nor dealt with the presence of interference from ambient environmental cues in real-world settings for the sensory cues. Further, a majority of the prior studies assessed memory performance only over less than a day, with about 5-12% improvement in post-nap memory performance on simple laboratory tasks (cf., Rudoy et al., 2009, Antony et al., 2012), and employed open-loop fixed-dose cueing during offline periods. Moreover, TMR with sensory cues may not be scalable to the multitude of experiences over one’s lifetime. Our study overcomes these limitations by investigating the effects of non-sensory transcranial electrical stimulation (tES) for TMR of real-world episodic memories with one-shot acquisition, and then assessing recall over a 48 hour period.

Prior work on non-sensory cueing showed that transcranial magnetic stimulation can reactivate the experience of a visual stimulus after repeated pairing with the stimulus (Liao et al., 2013). And transcranial alternating current stimulation (tACS) of left dorsolateral prefrontal cortex at a given frequency (60 or 90 Hz) during encoding can boost subsequent performance for old vs. new recognition of learned words when reapplied at the same frequency during either retrieval (Javadi et al., 2017) or slow-wave sleep (Crowley & Javadi, 2019). Employing multi-site intracranial recordings in non-human primates, we showed that transcranial direct current stimulation (tDCS) of prefrontal cortex alters functional connectivity between brain areas in a frequency-specific manner. Further, modulated inter-areal coherence in gamma frequencies accounted for improved learning speed on an associative memory task, without any changes in single-unit and multi-unit firing rates (Krause et al., 2017). Recently, we showed that tACS can consistently entrain the spiking activity of single neurons in deep structures in a spatially-localized and frequency-specific manner (Krause et al., 2019). Building on these prior results, we hypothesized that unique spatiotemporal amplitude-modulated patterns (STAMPs) of tES could potentially alter the functional connectivity as well as the temporal structure of spiking activity within the brain in unique ways and thereby be leveraged for TMR in more potent ways than sensory cues. In particular, we investigated whether such unique STAMPs of tES could be used to tag specific naturalistic episodes during one-shot viewing in immersive virtual reality and subsequently cue them during sleep to boost their memory recall over 48 hours in a targeted manner. The overarching goal of this study is to assess if coarser, non-invasive stimulation is sufficient to effectively modulate complex human memories.

## Methods

We computed a library of 256 STAMPs over a 32-channel array on the scalp (see Fig. S1) using gradient descent optimization for a realistic 3D adult human head template (see Supplementary Information). We employed immersive virtual reality (VR) to investigate the modulation of human episodic memory with ecological validity. Subjects were able to freely look around a fully rendered 3D environment on the HTC Vive^®^ platform. Their virtual vantage point was situated on a balcony across from an apartment building set in a nondescript Middle Eastern town (see Movie S1). The task was to actively surveil the inhabitants of the building and the passers-by so that they are able to later recall the events. We designed 28 episodes in this simulated environment, each about a minute long, and divided them into two groups of 14 episodes each (see Supplementary Information). To make the viewing more active and increase motivation, subjects were instructed to respond to salient events in each episode as they happened by orienting a reticle in the head-mounted display (HMD) towards those events and taking pictures with a virtual camera triggered by a hand-held controller.

The experiment was conducted over the course of seven days and included five nights in our sleep laboratory that comprised an acclimation night and four experimental nights (see Fig. 1A). Two experimental nights followed the acclimation night, whereas the other two took place about 8.25 days (standard deviation = 4.92 days) later. N=24 subjects were randomly assigned to one of four conditions in a within-subjects, counterbalanced, single-blind design based on which episode group (‘A’, ‘B’) and which stimulation condition (‘Active’, ‘Sham’) were employed in the first week. For the Active stimulation condition, each of the 14 episodes were stimulated with a unique STAMP once during viewing (see Fig. 1B). The pairing between STAMPs and episodes was arbitrary and randomly chosen for each subject. Only half of the STAMPs employed for tagging were re-applied to temporally coincide with the UP states of SWOs through the subsequent two experimental nights (see Fig. 1C). The corresponding episodes were termed “Tag & Cue”, and the other half of the episodes were termed “Tag & No Cue”. For the “Sham” stimulation condition, the 14 episodes were neither tagged during waking nor cued during sleep. We employed tDCS STAMPs for 8 subjects (termed “Spatial”) and 40 Hz tACS STAMPs for the remaining 16 subjects (termed “Temporal”). Note that while the same STAMPs were employed during waking and sleep, the duration and number of applications differed markedly; each waking STAMP was applied during the single exposure of the associated episode, about a minute long, whereas each STAMP was applied multiple times (37 to 284 times) during sleep, each about 0.4167 to 1 s long. See Fig. 2 for an illustration of the STAMP intervention.

**Fig. 1.**
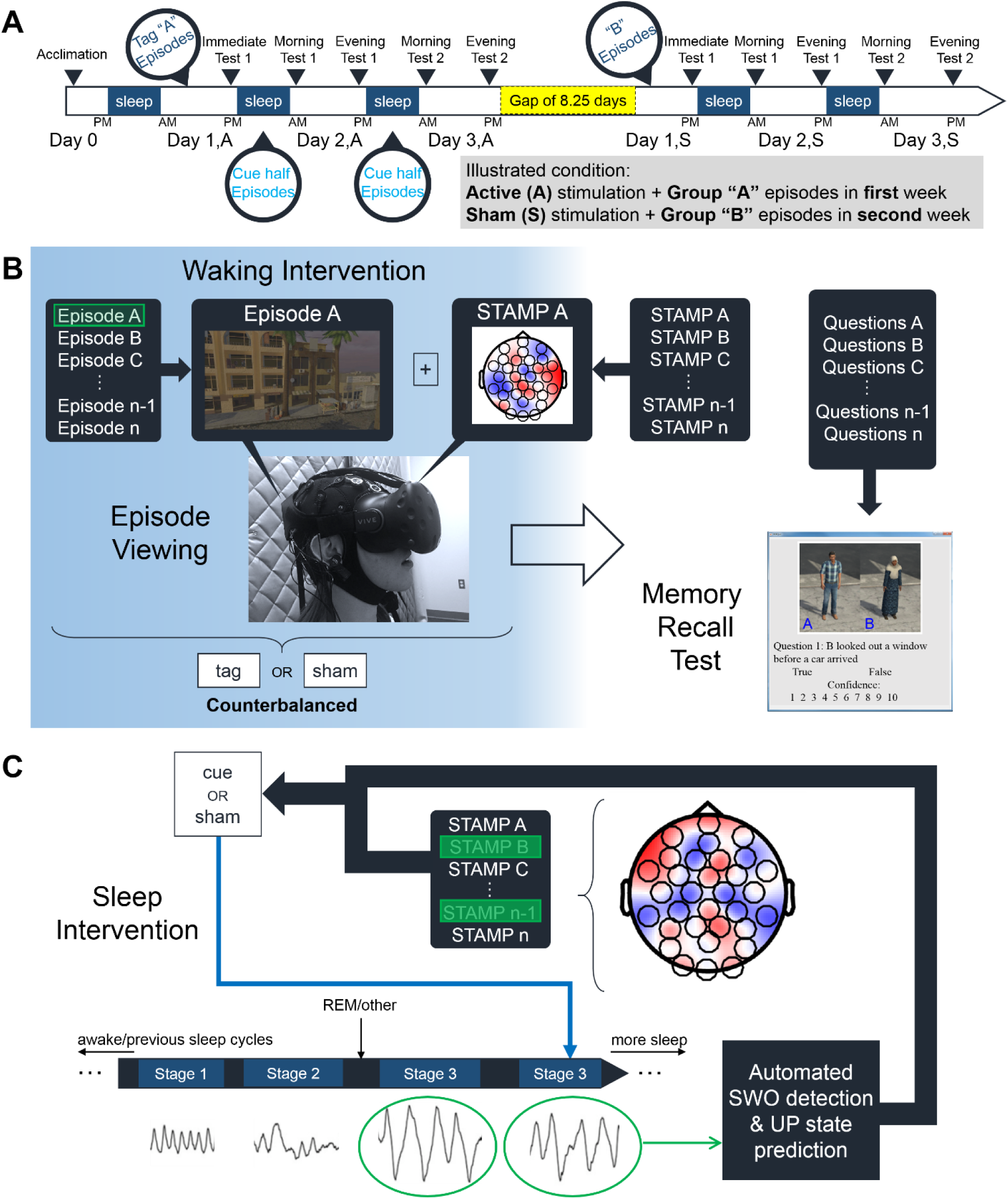
Experimental paradigm. (A) Subjects participated in a within-subjects counterbalanced study involving one acclimation night and a pair of two experimental nights. (B) Illustration of the experimental procedures during waking. (C) Illustration of the experimental procedures during sleep.

**Fig. 2.**
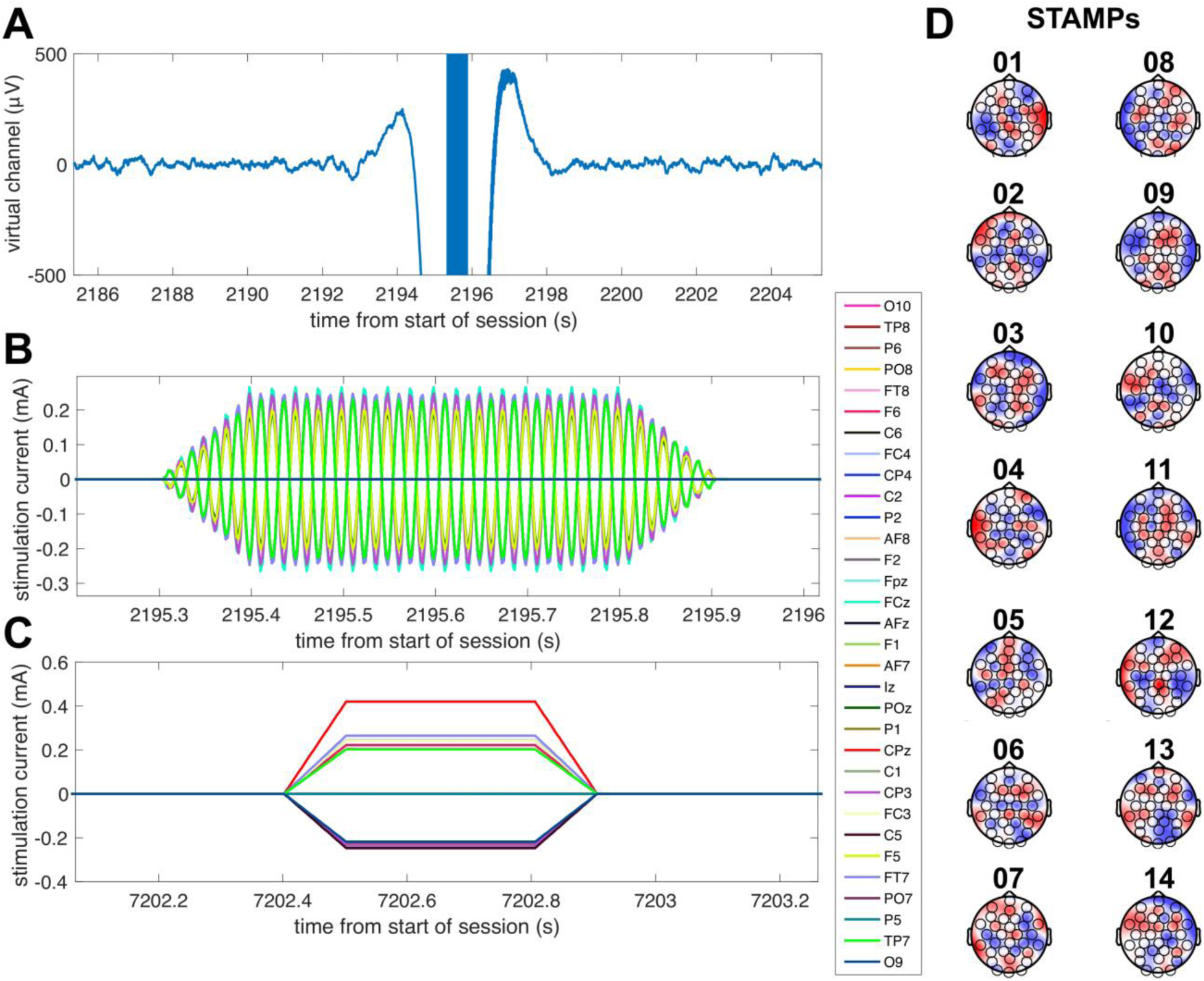
Illustration of STAMPs used in the experiment. (A) Time-locked plot of the EEG virtual channel, bandpass filtered between 0.5 and 50 Hz, as a tACS STAMP is applied during a predicted UP state of SWOs for a representative subject. Note the high-amplitude stimulation artifacts surrounding the application of the STAMP. (B) Time-locked plot of the applied currents for the tACS STAMP shown in (A), with 100 ms up and down ramps. (C) Plot of the applied currents for a representative tDCS STAMP, with 100 ms up and down ramps. (D) Scalp topographies of the amplitudes of currents for the 14 STAMPs that were used in the experiment to tag episodic memories.

For each stimulation condition, memory performance was assessed in five test sessions over the course of 48 hours following initial viewing of the episodes. In each test session, subjects determined the veracity of two declarative statements for each of the 14 episodes (see Fig. 1B and Supplementary Information), and also rated their confidence in the correctness of their response for each statement on a scale from 1 to 10. Textual prompts and pictures of characters were the only cues available to recall the pertinent episodes. Subjects were instructed to respond as quickly as possible without sacrificing accuracy. The correct/incorrect responses and the confidence ratings were used to generate Type-2 receiver operating characteristic (ROC) curves (Galvin et al., 2003; MacMillan & Creelman, 2005) for the three intervention types in each of the three days, which are segmented by the two experimental nights. The area under the Type-2 ROC curve served as the metric of episodic metamemory (see Fig. S2 and Supplementary Information). The metric is essentially the probability that a randomly chosen correct recall has a higher confidence rating than a randomly chosen incorrect recall. Our main hypotheses of interest were the performance for ‘Tag & Cue’ episodes would be better than that for either ‘Tag & No Cue’ or ‘Sham’ episodes following the sleep intervention with ‘Tag & Cue’ STAMPs.

## Results

We first validated whether our closed-loop algorithm applied STAMPs during the UP states of SWOs. Using electroencephalography (EEG) data from the Sham nights in order to avoid high-amplitude stimulation artifacts, the SWO phase at the start of each SW event was estimated from the time-locked event-related potential (ERP) of the EEG virtual channel. The distribution of mean SWO phases across subjects was found not to be significantly different from 0° (−2.18° ± 6°; v-test: v = 18.8, *p* < 0.001), which marks the start of UP states (see Fig. 3A). Accordingly, the grand average of ERPs time-locked to the start of the predicted UP states revealed a consistent transition from a DOWN state to an UP state at the appropriate time point (see Fig. 3B). We next analyzed the number of slow-wave (SW) events during sleep using a linear mixed-effects model with subject as a random factor and with fixed factors for stimulation condition (‘Active’, ‘Sham’), experimental night (‘Night 1’, ‘Night 2’), contrast coded factors comparing sleep stage N2 to average of all other stages and N3 to the average of all other stages, and all possible interactions among them (see Fig. 3C). There were significantly more SW events during N2 (t(369.3) = 6.673, *p* = 9.18e-11) and N3 (t(369.3) = 3.903, *p* = 0.000113) stages compared to all other stages. All other effects were not significant. These results verify the accuracy of automated SWO detection and UP state prediction for the closed-loop application of STAMPs.

**Fig. 3.**
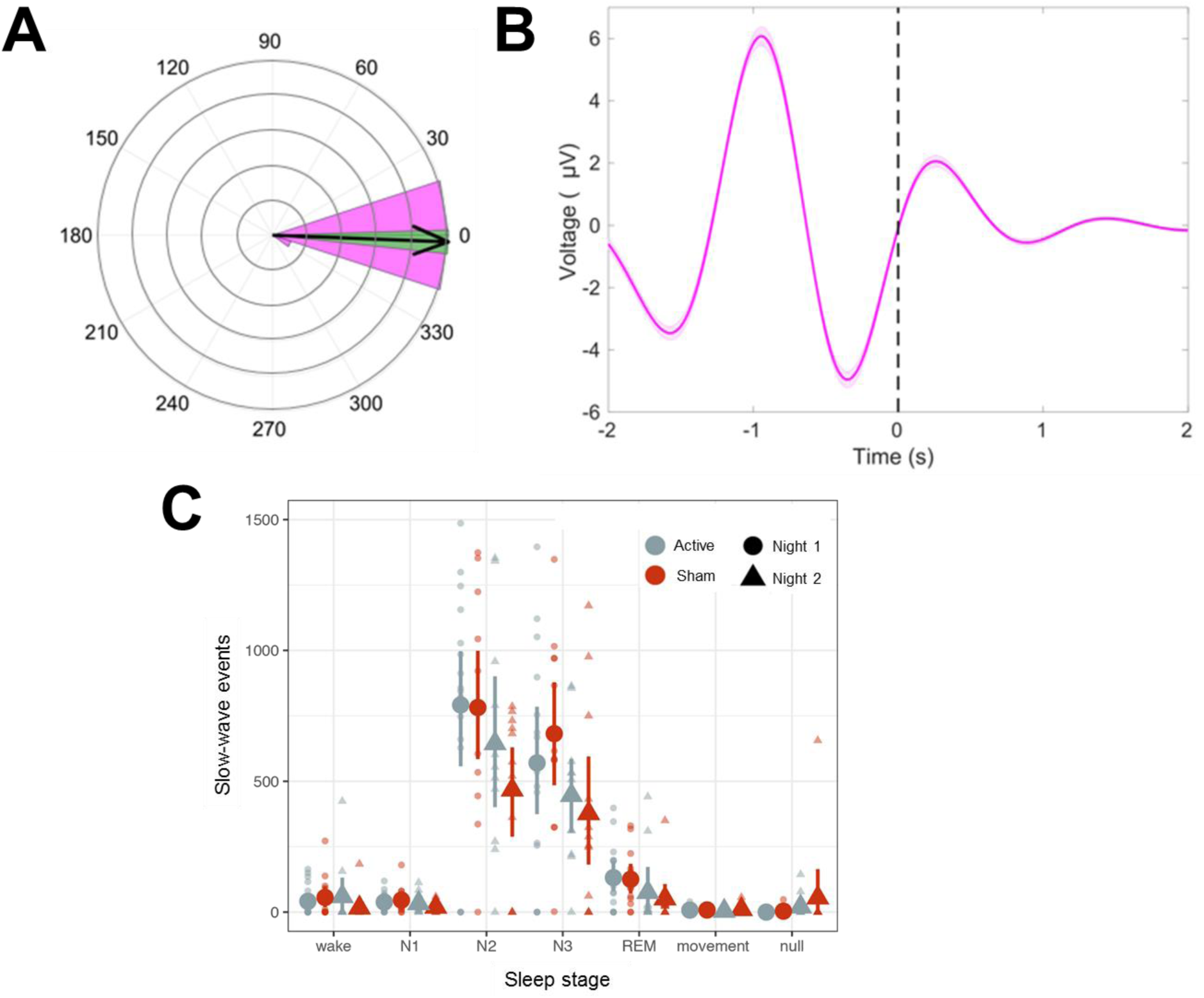
Validation of closed-loop application of STAMPs during UP states of SWOs in NREM sleep stages 2 and 3. (A) Grand average of the ERP of the virtual channel (bandpass filtered between 0.5 and 1.2 Hz) time-locked to the start of predicted UP states from the Sham nights across subjects. Shaded area shows the 95% CI. (B) Polar histogram of average phase values at the start of predicted UP states from the Sham nights across subjects (the pink shaded region), with the arrow marking the grand average. Green shaded region shows the 95% CI. (C) Distribution of overnight SW events in various sleep stages as determined by an expert human scorer. X-axis shows sleep stage categories including unstageable epochs labeled ‘null’. SW events in each stage are counted within subject for each stimulation condition (‘Active’ and ‘Sham’) and experimental night (‘Night 1’ and ‘Night 2’), shown in small transparent markers. The group mean and 95% confidence interval are shown on the large markers for each condition x night.

For behavior, we first examined absolute accuracy scores using a linear mixed-effects model with subject as a random factor. Fixed effects included intervention type (‘Tag & Cue’, ‘Tag & No Cue’, ‘Sham’), day (‘Day 1’, ‘Day 2’, ‘Day 3’), the interaction of intervention type and day, and covariates of STAMP type (‘Temporal’, ‘Spatial’), stimulation condition order (‘Active First’, ‘Sham First’), and episode group (‘Group A’, ‘Group B’). We did not find any significant effects (all *p*’s > 0.34), which suggests that STAMPs did not selectively modulate the absolute accuracy of memory recall as such.

We next analyzed the metamemory scores (see Fig. 4A) with a linear mixed-effects model. Fixed and random effects were same as above. Unlike absolute accuracy, metamemory significantly differed across days (F(2,192) = 4.8475, *p* = 0.008834), and there was a marginally significant interaction between intervention type and day (F(4,192) = 2.1201, *p* = 0.079842). All other effects were not significant. Based on these results, we ran follow-up linear mixed-effects models for each day separately with intervention type and subject as fixed and random factors, respectively. For Days 1 and 2, there were no significant effects. However, for Day 3, we found a significant main effect of intervention type (F(2,48) = 6.6238, *p* = 0.002882). Thus, the application of STAMPs during the exposure of ‘Tag & Cue’ and ‘Tag & No Cue’ episodes did not modulate their memory encoding as such, but the metamemory scores differed significantly across the intervention types by Day 3 following the application of ‘Tag & Cue’ STAMPs during two consecutive nights.

**Fig. 4.**
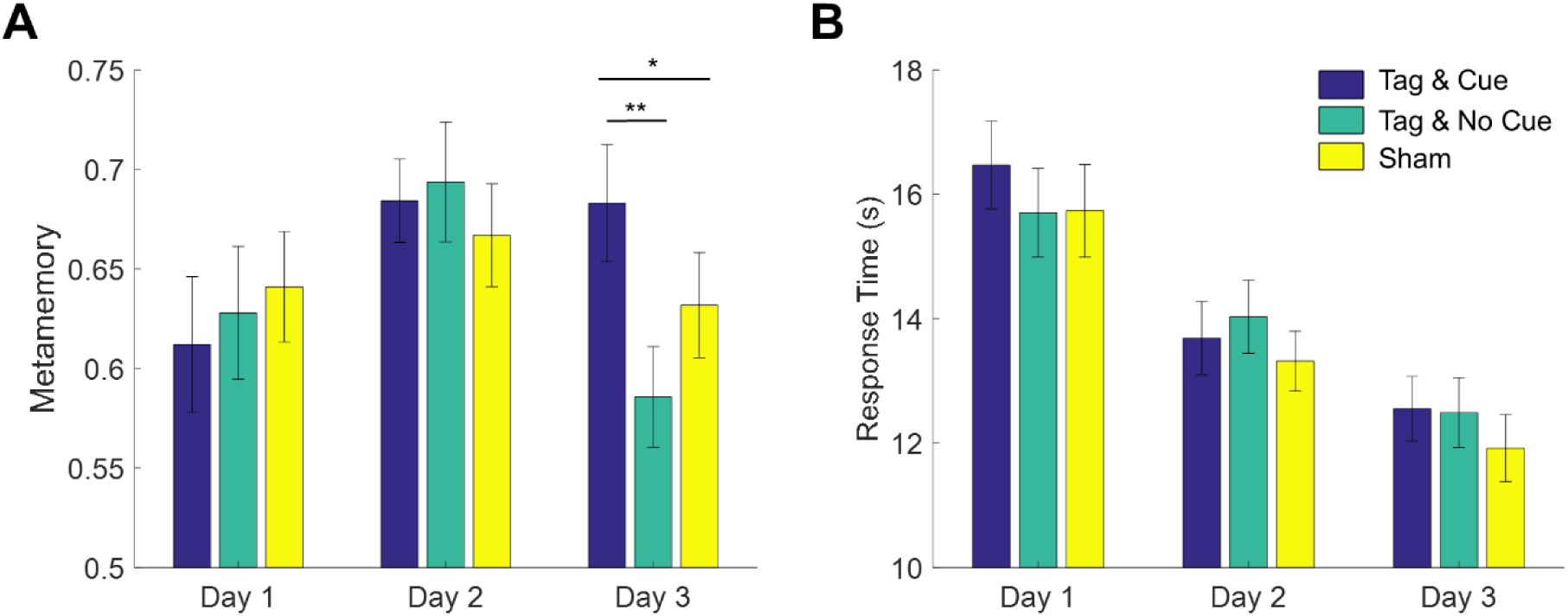
STAMP tagging and cueing enhances episodic metamemory without a speed-accuracy trade-off. (A) Metamemory performance on the episodic memory task is reported across the three days of testing. Blue bars show the metamemory scores for ‘Tag & Cue’ episodes, green for ‘Tag & No Cue’ episodes, and yellow for ‘Sham’ episodes. Metamemory scores for ‘Tag & Cue’ episodes are significantly higher than those for ‘Tag & No Cue’ by 19.43% (*p* = 3.7916e-3, corrected) or ‘Sham’ episodes by 10.01% (*p* = 0.04793, corrected) on Day 3. (B) Response time (RT) decreased across the three days of testing, but there were no significant differences among the intervention types. Error bars represent +/-1 SEM (N=24).

Given the significant main effect of intervention type on Day 3, we conducted two-tailed paired-sample t-tests on metamemory scores from Day 3. ‘Tag & Cue’ metamemory scores were significantly greater than both ‘Tag & No Cue’ (t(23) = 3.5069, adjusted *p* = 3.7916e-3, Hohm-Bonferroni method for 2 comparisons) and ‘Sham’ (t(23) = 2.0894, adjusted *p* = 0.04793, Hohm-Bonferroni method for 2 comparisons) metamemory scores. Thus, the application of STAMPs during SWOs in the two nights following one-shot viewing led to specific enhancement of metamemory of episodes that were both tagged and cued.

To understand the primary result better, we also analyzed response times (RTs) using a linear mixed-effects model with subject as a random factor (see Fig. 4B). Fixed effects included intervention type, day, and the interaction of intervention type and day. We only found a significant main effect of day (F(2,192) = 74.7045, *p* < 2e-16). This demonstrates that the benefit to metamemory from STAMP-based tagging and cueing was not simply due to a speed-accuracy trade-off, as subjects responded as quickly across all three intervention types (including on Day 3).

The immediate pre-sleep metamemory and the number of STAMP applications in each Active night varied widely across the subjects, with the latter depending on the number of SW events detected. We therefore analyzed the effects and interactions of these covariate variables on the immediate post-sleep metamemory for ‘Tag & Cue’ and ‘Tag & No Cue’ episodes within the Active stimulation condition using a linear mixed-effects model with subject as a random factor (see Fig. 5). Fixed effects of this model included categorical variables of intervention type (‘Tag & Cue’, ‘Tag & No Cue’) and experimental night (‘Night 1’, ‘Night 2’), continuous variables of pre-sleep metamemory and the number of STAMP applications, and all possible interactions among them. We found a marginal main effect of STAMP count (F(1,95.663) = 2.8666, *p* = 0.0936898), significant main effects of intervention type (F(1,83.846) = 13.2344, *p* = 0.0004738) and pre-sleep metamemory (F(1,94.743) = 4.4279, *p* = 0.0380035), a marginal two-way interaction of pre-sleep metamemory and STAMP count (F(1,93.544) = 3.8625, *p* = 0.0523440), significant two-way interactions of intervention type and STAMP count (F(1,82.014) = 7.6007, *p* = 0.0071894) and of intervention type and pre-sleep metamemory (F(1,84.598) = 11.4234, *p* = 0.0010991), and a significant three-way interaction of intervention type, pre-sleep metamemory, and STAMP count (F(1,82.829) = 7.0743, *p* = 0.0093864). All other effects were not significant. In particular, post-sleep metamemory was not modulated by experimental night.

**Fig. 5.**
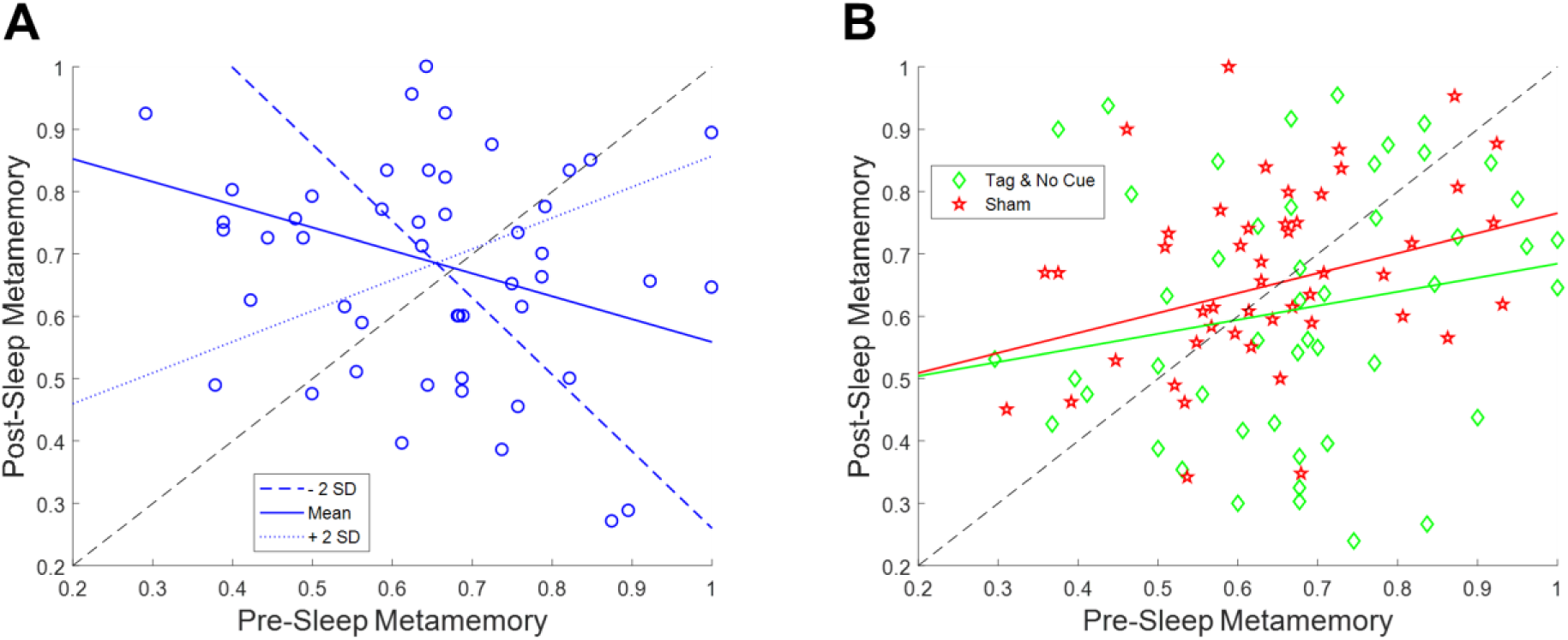
The effect of STAMP tagging and cueing on metamemory is modulated by an interaction between pre-sleep metamemory and the number of STAMP applications. (A) Post-sleep metamemory for ‘Tag & Cue’ episodes depends on the interaction between the covariates of pre-sleep metamemory and STAMP count. The three lines correspond to the linear mixed-effects model predictions for different STAMP counts (ranging from two standard deviations below the mean (blue) to two standard deviations above the mean (red). (B) The interactions of pre-sleep metamemory with STAMP count for ‘Tag & No Cue’ episodes, and with the number of SW events for ‘Sham’ episodes, are not significant. The markers in either panel correspond to data from individual subjects from both experimental nights.

Based on these results, we ran follow-up linear mixed-effects models for ‘Tag & Cue’ and ‘Tag & No Cue’ episodes separately with subject as a random factor and with pre-sleep metamemory, STAMP count, and the interaction of pre-sleep metamemory and STAMP count as fixed effects. There were no significant effects for ‘Tag & No Cue’ episodes. However, for ‘Tag & Cue’ episodes, we found significant main effects of both pre-sleep metamemory (F(1,45.954) = 9.1316, *p* = 0.004097) and STAMP count (F(1,43.033) = 7.4571, *p* = 0.009122), and a significant interaction of pre-sleep metamemory and STAMP count (F(1,41.848) = 7.7548, *p* = 0.008011). Given that the application of STAMPs coincided with the UP states of SWOs, we considered if the effects involving the STAMP count for the ‘Tag & Cue’ episodes were confounded by the number of concomitant SW events. To resolve this, we analyzed post-sleep metamemory for the ‘Sham’ episodes using a linear mixed-effects model with subject as a random factor and with continuous variables of pre-sleep metamemory and the number of SW events, a categorical variable of experimental night (‘Night 1’, ‘Night 2’), and all possible interactions among them as fixed effects. We did not find any significant effects (see Fig. 5B). These results provide further evidence for the specific modulation of episodic memories that were both tagged during waking as well as cued during sleep.

Across all the intervention types, sleep had a normalizing effect on metamemory, exhibiting an increase in subjects with weak pre-sleep performance and a decrease in those with strong pre-sleep performance. The application of STAMPs during sleep exaggerated this effect for ‘Tag & Cue’ episodes with a bigger overall overnight increase in metamemory for subjects with weak pre-sleep metamemory. But subjects with weak pre-sleep metamemory who received more than the mean dose of STAMP cueing during sleep had a lower overnight increase in metamemory than those who received less than the mean dose. On the other hand, subjects with strong pre-sleep metamemory who received more than the mean dose of STAMP cueing experienced less of an overnight decline in metamemory (see Fig. 5).

Finally, we investigated the neurophysiological effects of ‘Tag & Cue’ STAMPs during Night 2 owing to significant differences between ‘Tag & Cue’ episode and other intervention types on Day 3. First, we contrasted average post-SW event changes in spectral power between Active and Sham Night 2 using cluster-based permutation tests (see Supplementary Information) for five frequency bands (SW: 0.5-1.2 Hz, delta: 1-4 Hz, theta: 4-8 Hz, slow-spindle: 9-12 Hz, fast-spindle: 12-15 Hz), which are of relevance to memory consolidation (Mölle et al., 2011; Cox et al., 2014). Note high-amplitude stimulation artifacts preclude the inspection of EEG data during the application of STAMPs and up to 3 s following their offset. In short, significant channels from paired-samples comparisons of spectral power changes between Active and Sham Night 2 from 3 to 10 s following the offset of SW events were spatiotemporally clustered for each frequency band. The significant of these clusters, as determined by a permutation test, were defined as the “contrast clusters”. A similar cluster analysis was then carried out for correlations of differences in overnight changes in metamemory between ‘Tag & Cue’ and ‘Sham’ episodes with the corresponding differences in spectral power changes following SW events to obtain “correlation clusters”. This analysis was restricted to the channel × time bins in the significant contrast clusters, which showed a significant average change in spectral power over the night. We were thus able to associate overnight metamemory changes with specific spectral power modulations induced by STAMPs.

The analysis revealed a significant contrast cluster only in the slow-spindle band (9-12 Hz), such that the average post-SW event change in spectral power was lower (and negative) in the Sham Night 2 compared to the Active Night 2 (see Fig. 6A). This cluster temporally extended from 6.18 to 6.7 s relative to the offset of SW events, and had a Bonferroni-corrected clusterwise *P* value of 0.025 (*p* = 0.005, uncorrected). Scalp topography of t-values for the spectral power changes in the cluster indicated an early distribution over pre-frontal, left frontal, and left temporal areas, which then widened to include occipital and parietal regions (see Fig. 6B). Next, we correlated the differences in overnight metamemory changes between ‘Tag & Cue’ and ‘Sham’ episodes with the differences in spectral power changes within this contrast cluster between Active and Sham Night 2. We found a positive correlation cluster within the slow-spindle band between 6.56 and 6.64 s relative to the offset of SW events, with a clusterwise *P* value of 0.013 (see Fig. 6C), such that the stronger the average STAMP-induced increase in slow-spindle power had been, the higher was the overnight improvement in metamemory (see Fig. 6D). Scalp topography of the summed t-values for the spectral power changes in this correlation cluster indicated a concentration on the left temporal region.

**Fig. 6.**
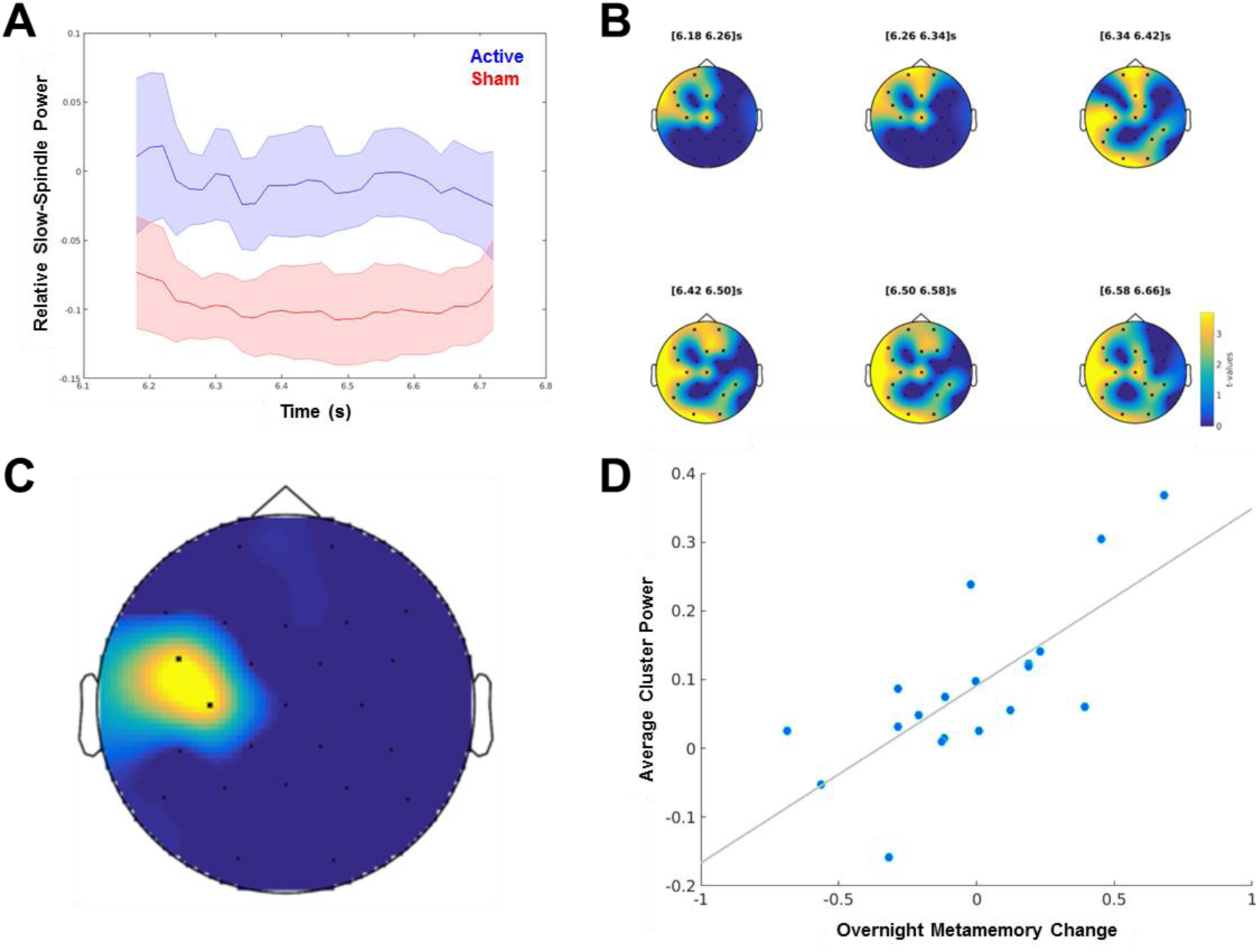
Neurophysiological and behavioral effects of STAMPs for Night 2. (A) Dynamics of spectral power changes for the slow-spindle (9-12 Hz) contrast cluster, relative to the offset of SW events, showing significant differences between Active and Sham Night 2. (B) Scalp topography of t-values for the slow-spindle contrast cluster (clusterwise *p* = 0.005) at six time points within the significant window (6.18 to 6.7 s following the offset of SW events). EEG channels participating in the cluster are marked by asterisks. (C) Scalp topography of summed t-values for the slow-spindle correlation cluster (clusterwise *p* = 0.013) from significant time bins. (D) Correlation of the differences in average cluster power from significant correlation cluster bins between Active and Sham Night 2 and the differences in overnight metamemory change between ‘Tag & Cue’ and ‘Sham’ episodes. Dots correspond to data from individual subjects (N=19).

## Discussion

We have demonstrated a novel form of tES in healthy humans that can boost the metamemory of specific naturalistic episodes that were viewed only once in immersive virtual reality. In particular, we discovered that unique spatiotemporal patterns (namely, STAMPs) of tES applied during SWOs can not only boost but also impair the metamemory of individual episodes depending on an interaction between pre-sleep metamemory and the number of STAMP applications through the night. Further, we found that post-stimulation increases in slow-spindle (9-12 Hz) power for left temporal areas in scalp EEG serve as a STAMP-induced biomarker of overnight metamemory improvements.

Though we found that STAMPs did not modulate the absolute accuracy of memory recall, we are not able to draw any conclusions regarding whether STAMPs can instead modulate bias-free measures of episodic memory such as those that use Type-1 ROC curve analysis of confidence ratings on a scale from “Definitely False” to “Definitely True” for the declarative statements about the episodes (Macmillan & Creelman, 2005; Mickes et al., 2012). Our experimental design only supports Type-2 ROC curve analysis (which measures metamemory) owing to the two-level responses – one for whether a declarative statement was “True” or “False”, and the other to rate the correctness of the response on a scale from “Least Confident” to “Most Confident”. Though pseudo confidence rating distributions for Type-1 true positives and false positives can be theoretically constructed by spanning the range from “Definitely False” to “Definitely True” using highest-confidence “False” and “True” responses, respectively (Gombos et al., 2012), the subjects’ Type-1 responses can only be merely predicted (with certain assumptions) rather than empirically measured (Galvin et al., 2003).

While it remains an open question if STAMPs also modulate episodic memory in a targeted manner, previous studies have suggested that memory and metamemory are likely correlated (Sacher et al., 2009; Yacoby et al., 2015). Broadly speaking, it has been proposed that metamemory is driven by either the familiarity of the recall cue, or the accessibility of any available pertinent information including that retrieved from memory in response to the cue (Koriat & Levy-Sadot, 2001). In either case, judgments about memories likely leverage the same neural representations of memory traces as the memory recall itself (Fleming & Dolan, 2012). The interaction between pre-sleep metamemory and STAMP count during sleep for ‘Tag & Cue’ episodes on post-sleep metamemory can be understood with the theory of complementary learning systems (McClelland et al., 1995). According to this theory, episodic experiences are rapidly encoded in the hippocampus during waking for the short term. Before the hippocampal memory traces fade out, episodic memories are consolidated into long-term storage in the slow-learning neocortex during sleep through replays of pertinent neural activity patterns. Subjects with weak metamemory prior to sleep benefit from STAMP-based cueing because the sequential structure of the episodes can be strengthened in the hippocampus as well as consolidated in the cortex. We speculate that an excessive number of STAMPs can, however, roll back this benefit by learning remote, higher-order links between events within the episodes. Subjects with strong metamemory prior to sleep benefit do not benefit from STAMP cueing as such due to the same reason. So, the prescription for boosting the metamemory of episodic experiences is to apply an optimal number of STAMPs during sleep for subjects with weaker pre-sleep metamemory, and to not intervene for subjects with stronger pre-sleep metamemory (cf., Schapiro et al., 2018).

The effect of STAMPs on overnight metamemory improvements was correlated with an increase in slow-spindle (9-12 Hz) power for left temporal areas following STAMPs during sleep. Spindles are demonstrably a critical component of sleep-dependent memory consolidation (Fogel & Smith, 2011), as shown in rodents (Eschenko et al., 2006) as well as humans (Cox et al., 2012). Recent optogenetic work in rats demonstrated a causal role for spindles in coupling SWOs and ripples for effective consolidation (Latchoumane et al., 2017). Spindles reverberate in circular wave-like patterns across temporal, parietal, and frontal regions repeatedly throughout the night, regulating the process of memory re-organization over time and space (Muller et al., 2016). To the best of our knowledge, there have been no studies on the hemispheric differences in the sleep-dependent consolidation of episodic memory and/or metamemory. In this regard, our results suggest that the left temporal lobe may play a key role in boosting the consolidation of episodic metamemory.

In summary, we have developed a novel non-sensory non-invasive method to tag naturalistic episodic experiences in one shot and cue them during offline periods to boost their metamemory in a targeted manner. New sleep studies are in order that aim to maximize the benefits of STAMPs during sleep by regulating the number of STAMP applications based on pre-sleep metamemory, as well as the intensity and frequency of particular STAMPs based on post-stimulation biomarker of metamemory. Our results suggest that, unlike relatively localized brain circuits responsible for regulating mood (Rao et al., 2018) and movement (Follett et al., 2010), episodic memories are processed by a much more widespread network of brain areas. We believe our study will pave the way for next-generation transcranial brain-machine interfaces that can dramatically boost learning and memory in healthy humans for real-world tasks. Such a non-invasive approach can also potentially benefit a majority of patients with learning and memory deficits at much lower cost and risk than required for implanting intracranial electrode arrays. It is also possible to dramatically enhance the efficacy of exposure behavioral therapy using a combination of immersive virtual reality and STAMP-based tagging and cueing for the treatment of post-traumatic stress disorder.

## Acknowledgements

We would like to thank Neuroelectrics, Inc. for contributions to the generation of STAMPs and for building the prototype StarStim 64 device, Dr. Albert Rizzo and Dr. John Wixted for contributions to the design of the behavioral paradigm, Aashish Patel for contributions to the behavioral analysis, and Dr. Mark Gluck and Dr. Penelope Lewis for useful discussions through the course of this study. This material is based upon work supported by the DARPA and the Army Research Office under Contract No. W911NF-16-C-0018 (RAM Replay program). The views, opinions and/or findings expressed are those of the author and should not be interpreted as representing the official views or policies of the Department of Defense or the U.S. Government.

## Author contributions

Designed the experiment: P.K.P., M.D.H.; Developed the behavioral task: P.K.P., S.W.S., N.A.K., S.M.R., I.L., A.H., J.C., M.D.H.; Ran the experiment: A.P.J., B.R., N.B.B., T.S.M., V.P.C.; Performed the analysis: P.K.P., R.J.H., N.A.K.; Wrote the paper: P.K.P., M.D.H.

## Supplementary Information

### Materials and Methods

#### STAMPs

Each STAMP is defined as an array of currents across 32 stimulation electrodes located in the 64-channel layout as per the international 10-10 system (see Fig. S1). A library of 256 spatial stimulation patterns was computed based on the criterion that the induced electric fields in the 3D brain volume of a standard adult human head template are as mutually orthogonal as possible. Electric field space was defined in the cortical mesh that had 190521 dimensions. Gradient descent optimization was used to minimize the norm of the cross-correlation function for electric fields across the library. Given that both positive and negative correlations were penalized, the computed spatial pattern solutions can be used for either tDCS or tACS STAMPs. Other constraints included the total injected current set to 2.5 mA, with maximum 1.5 mA and minimal 150 µA current at any electrode (to avoid impedance issues). Different initializations of the gradient descent search, in terms of the number of starting non-zero current electrodes, yielded STAMP solutions with different sparseness amounts. Solutions using more initial non-zero current electrodes led to STAMP sets with lower overall cross-correlation and lower overall currents across the montages. STAMPs used in the current study were solved based on initialization of 18 non-zero current electrodes. Of the 256 computed STAMPs, 14 were randomly chosen for usage in the current study. See Fig. 2D for the scalp topography of these 14 STAMPs.

#### Subjects

Subjects were 18-40 years of age, used English as a first language, completed high school, and had no history of head injury with loss of consciousness for longer than five minutes. They were right-handed according to the Edinburgh Handedness Inventory (Oldfield, 1971), had no history of neurological or psychiatric disorder, had no history of alcohol or drug abuse, were non-smoking, had no excessive alcohol or caffeine consumption, were not currently taking any medication significantly affecting the central nervous system, had no implanted metal, had no sensitivity or allergy to latex, had good or corrected hearing and vision, and reported no sleep disturbances. Women who were pregnant, or thought they may be, were also excluded.

A total of 30 healthy subjects completed the experiment. They were recruited using flyers placed around campus of the University of New Mexico and surrounding community, and received monetary compensation upon completion of the study. Of these, 6 subjects were excluded from the analyses due to either equipment failure during an Active night, or non-compliance in following task instructions. All subjects provided signed informed consent to participate in the study, which was approved by the Chesapeake Institutional Review Board. The remaining N=24 subjects (15 female) had a mean age of 23.96 years with a standard deviation of 6.08 years.

#### Behavioral Paradigm

We employed immersive virtual reality (VR) to produce simulated realistic environments for the purpose of investigating the modulation of human episodic memories with STAMPs. VR-derived results have greater predictive validity and relevance for real-world applications when compared to results from standard training and testing tools on a personal computer. In addition, on a more pragmatic level, rather than relying on costly physical mock-ups of functional environments, VR offers the option to produce and distribute identical “standard” simulation environments. Within such digital assessment scenarios, normative data can be accumulated for performance comparisons in a systematic manner.

As mentioned in the Introduction section, we designed 28 distinct memorable episodes in the VR. Each series of events in an episode generally centered around two main characters, often with one or two less involved characters. 10 declarative statements were composed for each episode to test the ability of the subjects to recall facts about the experienced events. The veracity of these statements can be determined by remembering the underlying story (i.e., the sequence of events) and which characters were involved. Movie S1 provides an illustrative episode (namely, “Fire Response”).

In preparation for the experiment, the episodes and questions (i.e., declarative statements) were gradually improved using feedback from 12 pilot subjects. They watched one episode at a time and immediately rated the difficulty of each of the 10 questions on a scale from 1 (easy) to 10 (difficult), and the overall memorability of the episode on a scale from 1 (least memorable) to 10 (most memorable). Average difficultly rating in the subjects’ responses tended to covary with the accuracy of their responses across the episodes.

We employed an iterative process that was aimed at subjectively equalizing the overall memorability and question difficulty across the episodes. Each iteration involved getting responses from four pilot subjects, after which the less memorable episodes were altered to increase memorability by adding more salient events. Questions that were answered incorrectly and rated as difficult, across the subjects, were made easier, and those that were answered correctly and rated as easy were increased in difficulty. After three iterations, the 28 episodes were of similar memorability, and the questions were of similar difficulty. Using data from one final cohort of four pilot subjects, the set of the 28 finalized episodes was curated into four subgroups with the constraints that the average difficulty of the questions and the average memorability of the episodes in each subgroup were similar, and the themes of the events occurring in the episodes were roughly evenly distributed. Two groups of 14 episodes each (namely, A and B) were randomly created by choosing among these four subgroups. Episodes were given names but were not presented to the subjects and were used only for reference by the experimenters (see Table S1). The 10 finalized questions for each episode were then randomly divided into five pairs, with each of the pairs assigned to one of the five test sessions (namely, quizzes) that were administered to the subjects.

For each stimulation condition, the quiz sessions were administered immediately following the episode viewing (termed “Day 1”), the morning after the first night (termed “Day 2 morning”), the evening after the first night (termed “Day 2 evening”), the morning after the second night (termed “Day 3 morning”), and the evening after the second night (termed “Day 3 evening”). Each quiz session was composed of two questions per episode for a total of 28 questions per session. At the end of the five quiz sessions, subjects had answered 10 questions per episode for a total of 140 questions. The presentation order of questions within a session were pre-randomized such that a subject would answer one question for all 14 episodes before they would see the second question for a given episode. The pre-randomization was performed at the experiment outset such that every subject received questions in the same order for a given episode group.

Quizzes were conducted via a custom graphical user interface (GUI) coded in MATLAB and administered in the sleep laboratory on the same computer used for the VR episode viewing. The GUI was a single application window that cycled through three sections as the subject answered questions. The first section the subject encountered for an individual question contained a textual prompt that described the episode this question pertained to and a button labeled “View Question” that the subject pressed when they were ready to proceed to the next section. The second section was divided into two panels. The first panel contained the same textual prompt from the previous screen, a list of the two to four characters from the episode presented in a pre-randomized order with a name (e.g., ‘Character B’) and a 99×142 pixel image in a neutral location and position, a question pertaining to a specific detail of the episode, and two radio buttons for “True” or “False” selection. The second panel contained 10 radio buttons for the subject to rate their confidence in the answer they provided in the first panel with 1 being “Least Confident” and 10 being “Most Confident”. There was no deadline for these responses. Finally a button labeled “Next Question” became active once the subject had answered the question and provided a confidence rating. This button led to a third screen that enforced a four second interval between questions and then automatically loaded the prompt screen of the next question. This process repeated until all questions for the session had been answered. No feedback on the accuracy of their answers was provided to the subjects at any time during the experiment. Subjects were instructed to respond as quickly as they can without sacrificing accuracy.

Data was automatically collected from the GUI during user interaction and saved for later analysis. Five data points were captured for each question: the True/False response, the confidence rating, the prompt screen viewing duration, the interval from the question screen presentation to the selection of a True/False response, and the interval from the question screen presentation to the selection of a confidence rating. Movie S2 demonstrates the GUI that was used to administer the memory recall tests.

#### Experimental Procedure

At the orientation session before the experiment began, subjects were invited to provide informed consent, and were given several questionnaires to assess various aspects of their personality and sleep habits, as well as to gather an IQ estimate. Following the questionnaires, head measurements were made (circumference, nasion to inion, and pre-auricular to pre-auricular) to fit a neoprene head cap. Subjects were next given a tour of the sleep laboratories and an explanation of the EEG/tES equipment and experimental procedures. Finally, each subject was issued a Fitbit wrist-worn biometric sensor (Dickinson et al., 2016) with instructions on how to correctly operate it to track sleep prior to their subsequent laboratory visits.

For the acclimation night, subjects arrived at the sleep laboratory by 17:00, and were prepped and fitted with a neoprene head cap to collect EEG data. An adapted version of Raven’s Progressive Matrices called Sandia Matrices was then administered (Matzen et al., 2010). Next, data was collected to calibrate biometrics (Patel et al., 2018) for use in a predictive computational model, including a breath count task to measure attentional lapses (Braboszcz and Delorme, 2011) that lasted 30 minutes, as well as a 3-back task to generate cognitive stress and mental fatigue (Hopstaken et al., 2015) that lasted about 21 minutes. Subjects could then relax in the laboratory until roughly 21:00, when they were prepped for PSG recording and tES during sleep. EEG electrode locations were digitized using Polhemus FASTRAK System (Polhemus, Inc.) for data analysis purposes as well as to measure how much the cap may have shifted during the subsequent sleep session. Subjects were instructed to lie down in a supine position at approximately 22:00, when biocalibrations were performed to help identify sources of noise in later EEG acquisition. This included EEG data collection of eyes open for 1 minute, closed for 1 minute, looking up, down, right, and left, blinking slowly 5 times, clenching the jaw, and finally moving into a comfortable sleeping position. Lights out for the subjects occurred between 22:00–23:00, and they slept for up to 8 uninterrupted hours before being awoken. During sleep, EEG data were monitored, and the closed-loop prediction algorithm was initiated when 4 minutes of continuous N2/N3 sleep was observed by a trained research assistant. Unrelated to the episodic memory task and STAMP-based intervention on the experimental nights, half of the subjects received closed-loop SW tACS (Ketz et al., 2018; Jones et al., 2018) on their acclimation night. This tACS condition was counterbalanced to the order of the stimulation conditions in the main experiment. Upon waking, subjects could use the restroom and were offered water and snacks. They filled out the Karolinska Sleep Diary (KSD; Åkerstedt et al., 1994) to assess subjective sleep quality. Next, they completed a 1-back task for about 21 minutes to assess alertness and were then disconnected from the EEG/tES hardware and released.

In the evening of the acclimation night, subjects were familiarized with the virtual reality environment and task procedures via a short practice session. During this session, subjects viewed one 3.5-minute long example episode in the virtual world and then completed a 10-question sample quiz on a personal computer. Both were guided by a trained research assistant who provided detailed instructions to the subject and answered any questions that arose.

For the first experimental night, subjects arrived at the laboratory at approximately 19:00, and were immediately set up for EEG data collection and STAMP stimulation. They first completed a brief baseline mood questionnaire to assess potential effects of STAMP stimulation on subjective mood. The mood questionnaire consisted of nine questions on a 0-5 Likert scale. Items included feelings of nervousness or excitement, tiredness, confusion, sadness, degree of frustration, dizziness, nausea, degree of physical pain or discomfort, and ability to pay attention. Next, we gave an introduction to the physical sensation questionnaire, which assessed subjective sensations due to stimulation.

The subjects then sat in front of the computer, put on the HTC Vive^®^ headset, and heard the task instructions. They viewed 14 episodes from either Group A or B, depending on their assignment, in a random order within the VR environment. They were instructed to take pictures of anything they deemed relevant to help them subsequently recall about the unfolding storyline. Following episode viewing, subjects rated three different types of sensations (itching, heat/burning, and tingling) on a 0-10 Likert scale, where 0 indicated no sensation at all and 10 indicated the most intense possible sensation. Any report of a seven or above would have resulted in immediate termination of the experiment, without penalty to the subject. No subjects were lost due to this reason.

Next, subjects completed a test (“Immediate Test”) to assess memory recall where they answered 28 true/false questions (two for each episode) as well as provided a confidence rating for each response. Following the quiz, subjects completed an exit mood questionnaire, identical to the entrance questionnaire, as well as a questionnaire assessing their strategy for the task. Then subjects were prepped for polysomnographic (PSG) recording during sleep at roughly 21:00. Lights out occurred between 22:00-23:00, and rise time was between 06:00-07:00. During sleep, a trained research assistant monitored EEG data and started the closed-loop stimulation algorithm at 4 minutes of continuous, visible N2/N3 sleep, which was then allowed to run through the remainder of the night. Subjects either received active (2.5 mA) or sham (no current) STAMPs for the entire duration of sleep. Stimulation was paused if the subject showed signs of waking and resumed after the subject was again in N2/N3 sleep. Upon waking, subjects filled out the Karolinska Sleep Diary (KSD; Åkerstedt et al., 1994) to assess subjective sleep quality. Next, they completed another test (“Morning Test 1”) to assess the effect of sleep consolidation on memory performance. They then filled out the strategy questionnaire, were disconnected from the EEG hardware, and released.

For the second experimental night, the procedures were similar to the first experimental night, with the following differences. Prior to EEG setup, subjects completed a Language History Questionnaire (LHQ). Subjects did not view the episodes from the previous evening again, but they completed another test (“Evening Test 1”) on them. Additionally, a new set of Sandia Matrices was administered. PSG setup and STAMP stimulation procedures were identical to the first experimental night. Upon waking, subjects completed another memory recall test (“Morning Test 2”). For the follow-up to the second experimental night, subjects arrived approximately 24 hours after their previous day arrival (19:00), were prepped for EEG data collection, and completed a final test (“Evening Test 2”) to assess the effect of sleep stimulation on more long-term memory retention.

After about 8.25 days (standard deviation = 4.92 days), subjects returned to the laboratory for their third and fourth experiment nights, succeeded by the final follow-up. The timeline and procedures were identical to the first and second experimental nights and their follow-up, the only differences being the group of episodes viewed in the VR (‘A’, ‘B’) and the stimulation condition (‘Active’, ‘Sham’) were opposite of their assignments for the first and second experimental nights. Upon completion of the follow-up to the fourth experimental night, a final exit questionnaire was administered to gather subjective ratings from subjects in terms of how they felt the intervention impacted their memory function in general. And they were debriefed, during which time they could ask questions about the nature of the experiment.

#### Waking Electroencephalographic (EEG) Data Collection

32-channel physiological data collection and simultaneous 32-channel stimulation were conducted using the StarStim64 device (Neuroelectrics, Inc.). The 64 electrodes were held in place using a neoprene head cap, according to the international 10-10 system (recording: *P7*, T7, *CP5*, *FC5*, *F7*, F3, *C3*, *P3*, *FC1*, *CP1*, *Pz*, PO4, *O2*, Oz, *O1*, PO3, *CP2*, *Cz*, *FC2*, *Fz*, AF3, *FP1*, *FP2,* AF4, *P8*, T8, *CP6*, *FC6*, *F8*, F4, *C4*, *P4;* stimulation: O10, TP8, P6, PO8, FT8, F6, C6, FC4, CP4, C2, P2, AF8, F2, FPz, FCz, AFz, F1, AF7, Iz, POz, P1, CPz, C1, CP3, FC3, C5, F5, FT7, PO7, P5, TP7, O9). Solidgeltrodes (NE028, Neuroelectrics, Inc.) and pistim electrodes (NE024, Neuroelectrics, Inc.) were used for physiological data collection and stimulation, respectively. EEG data was collected from 23 of these 32 sites (marked in italics above). The remaining electrodes (PO3, PO4, Oz, AF3, AF4, F3, F4, T7, T8) were repurposed to record electrocardiogram (ECG), electrooculogram (EOG), and electromyogram (EMG) to allow for the detection of artifacts and sleep stages. An ECG lead (PO3) was placed under the left collarbone, and both vertical (AF3) and horizontal (AF4) EOG were collected: one lead placed superior and lateral to the right outer canthus, and another lead inferior and lateral to the left outer canthus. The physiological data was sampled at 500 Hz. Common Mode Signal (CMS) and Driving Right Leg (DRL) reference electrodes (stricktrodes: NE025, Neuroelectrics, Inc.) were placed on the right preauricular. No online hardware filtering, except for line noise (60 Hz), was applied during collection.

#### Polysomnographic (PSG) Data Collection

For polysomnographic (PSG) data collection during sleep, the setup was nearly identical to wake, with a few exceptions. First, two EMG electrodes were placed on and under the chin in accordance with PSG recording guidelines set forth by the American Academy for Sleep Medicine (Berry et al., 2015) to help with sleep scoring. Second, data was collected from 25 EEG electrodes, of which C3, C4, O1, O2, F3, and F4 were used for sleep staging.

#### Waking STAMP Stimulation

For the Active stimulation condition, STAMP montages were delivered via the StarStim64 device (Neuroelectrics, Inc.) during the one-shot viewing of episodes in the VR. For all subjects, each episode was randomly assigned a unique STAMP from the set of 14 montages. A given STAMP was applied for the entire duration of the corresponding episode (about a minute long) with ramp up and ramp down times of 100 ms. Inter-episode interval was randomly sampled between 6 and 8 s.

#### Closed-Loop Stimulation During Slow-Wave Oscillations

Our closed-loop algorithm was automatically triggered through the whole night to transiently apply ‘Tag & Cue’ STAMPs during predicted UP states of SWOs. The algorithm first detects the presence of SWOs, which consist of low-frequency synchronized upward and downward deflections of EEG. It next attempts to match the stimulation frequency and phase with ongoing endogenous SWOs such that the maximal stimulation occurs at their UP states (positive half waves). For robust SWO detection, a virtual channel is computed by averaging 13 fronto-parieto-central EEG channels (Cz, FC1, FC2, CP1, CP2, Fz, C4, Pz, C3, F3, F4, P3, P4 in the international 10-10 system) to determine the overall synchronous activity of EEG recorded during sleep. The virtual channel allows the observation of moments of relatively high SW power, referred to as ‘SW events’, while averaging out activity of lesser magnitudes on individual channels unrelated to the pattern of SWOs. The included channels are stored in a running 5 s buffer. They undergo moving average subtraction with a 1 s window (to mean center the signals at 0 μV), and noisy channels exceeding 500 μV min-to-max amplitude across the 5 s are rejected before the virtual channel is computed. The buffer is updated with each discrete data fetch operation that gets the new latest data up to the point of data request. By the time the buffer is updated, there is a random transmission delay, which needs to be accounted for to plan and precisely time the stimulation in the near future.

The virtual channel data in the buffer is further processed to detect the presence of SWOs and predict the upcoming UP state. The algorithm applies a Fast Fourier Transform (FFT) to the buffered data to determine the power spectrum. Stimulation is planned when the ratio of the cumulative power in the SW band (0.5-1.2 Hz) is more than 30% of the total cumulative power from 0.1 to 250 Hz. If this SW relative power threshold is crossed, the data is bandpass filtered in the SW band with a second-order zero-lag Butterworth filter. The Hilbert transform is then applied to the filtered signal to obtain the analytic signal, and the phase of the analytic signal is shifted back by 90° to align it with the SWOs. Next a sine wave is fit to the imaginary component of this signal by optimizing the amplitude, offset, and phase parameter values and using the dominant frequency in the SW band from the power spectrum. The sine wave is then projected into the future, identifying the temporal targets that would synchronize STAMPs to the predicted endogenous SWOs. Throughout this process, the dynamic latency associated with data processing is timed using the system clock. Together with distributions of calibrated latencies for data fetch and stimulation commands (mean = 5 ms, standard deviation = 2 ms), which were measured offline, the algorithm estimates the correct time point to communicate with the hardware to initiate the stimulation. As an example, suppose that at a given moment the algorithm initiates data fetch to populate the buffer with the last five seconds of EEG data. The data then becomes available for processing a few ms (say, 6 ms) into the future based on sampling from the distribution for data fetch latency. Assume it then takes 100 ms for data processing to predict that the next UP state will occur 600 ms in the future from the starting time point. If it takes a few ms (say, 7 ms) to physically initiate stimulation based on sampling from the distribution for stimulation command latency, the algorithm will wait 487 ms (600 ms – 100 ms – 7 ms – 6 ms) after the EEG processing step to send the stimulation command to the device.

During Active nights, STAMPs assigned to ‘Tag & Cue’ episodes were administered through the sleep period to boost the probability of specific memory replays. In particular, these STAMPs were applied in closed loop to temporally match the predicted UP states of SWOs because memory replays in the cortex are associated with those states (Ji & Wilson, 2007). Sometimes, due to hardware and/or processing delays, the targeted start of the UP state stimulation was not possible. In these cases, the algorithm compared the current time to the (now deprecated) stimulation start time, and checked if at least 300 ms of UP state stimulation was still possible. If so, the stimulation was initiated immediately and continued through the remainder of the UP state. If this was not possible, the algorithm started stimulation at the next upcoming UP state based on further projection of the sine wave fit from the buffer. Once STAMP delivery was completed (i.e., after stimulation offset), the system idled for 3 s to avoid the collection of high-amplitude stimulation artifacts in the data buffer, then resumed the cycle of data update in the buffer in search of the next SW event, at which point another STAMP was administered, and so on. STAMPs had ramp up and ramp down times of 100 ms during sleep as well. A minimum interval of 8 s was imposed between two consecutive SW events. Further, the seven STAMPs for the ‘Tag & Cue’ episodes were always applied in a batch, with their order randomized across batches through the sleep period.

Thus, the closed-loop stimulation system was able to target UP states of the ongoing endogenous SWOs, while adapting parameters continuously in real time in order to minimize temporal inaccuracies due to hardware and/or processing delays. For the Sham stimulation condition, the same algorithm was applied to mark the potential stimulation times without any currents being actually applied.

#### Type-2 ROC Analysis

Metamemory was calculated empirically from correct and incorrect recalls and the corresponding confidence ratings using standard procedure (Galvin et al., 2003; Macmillan & Creelman, 2005; Fleming & Lau, 2014). In each test trial, the textual prompt and the pictures of relevant characters would help retrieve the corresponding episode. The declarative statement would then be matched with the retrieved episode to determine its veracity. The stronger the memory recall, the greater is the ability to correctly recall the events from the episode without confabulating any details. A correctly answered declarative statement (irrespective of the response category) was considered a Type-2 true positive, and an incorrectly answered declarative statement (irrespective of the response category) a Type-2 false positive. Relative frequency distributions of confidence ratings (on the scale from 1 to 10) with 10 bins were computed for Type-2 true positives (i.e., correct recalls) and for Type-2 false positives (i.e., error recalls) separately. Next, the cumulative distribution functions were calculated for each in reverse direction (from end to beginning). The data points of the cumulative distribution function for Type-2 true positives were plotted against the corresponding data points of the cumulative distribution function for Type-2 false positives to obtain the Type-2 ROC curve. As noted in the Introduction section, the area under the Type-2 ROC curve was employed as the metric of metamemory for the episodic memory recall (see Fig. S2).

#### Post-hoc Sleep EEG Analyses

Sleep EEG analyses were run on N=19 subjects, as the data from 5 subjects could not be used because of excessive artifacts in one or more experimental nights. For sleep staging, EEG data of electrodes C3, C4, O1, O2, Fp1, and Fp2 were bandpass filtered between 0.5 and 35 Hz, together with EMG data between 10 and 100 Hz. Each 30-second epoch was visually inspected by an experienced technician and assigned a stage of wake, N1, N2, N3/SWS, REM, or movement according to guidelines by the American Academy for Sleep Medicine (Berry et al., 2015). Time in each sleep stage was directly derived by summing up all epochs determined to belong to that sleep stage. Percent of time in a sleep stage out of total sleep time was calculated as the amount of time spent in the stage divided by the total amount of time spent asleep.

We validated the UP state prediction algorithm using the data from Sham nights. This avoided the high-amplitude stimulation artifacts in the data from Active nights. Epochs time-locked to the start of predicted UP states (−5 to +5 s) were extracted from the EEG virtual channel data and bandpass filtered between 0.5 and 1.2 Hz. Phase values at the time point of each start marker were calculated using the Hilbert transform. Mean phase values across trials were calculated for each subject and then submitted to a *v*-test to examine whether the average across subjects is different from 0° (Berens, 2009). As noted above, in some instances a timed SW event either did not initiate at the start of an UP state or was moved to the next UP state, which may have increased the variability of these SW phases across the night to some degree.

Sleep EEG data were analyzed with custom-built scripts implemented in Matlab R2016a (The MathWorks), utilizing FieldTrip (Oostenveld et al., 2011) and EEGLab (Delorme & Makeig, 2004) functions. Data were epoched into pre- and post-SW event windows: pre-SW event windows captured -6.4 to -0 s before the onset of SW events, and post-SW event windows captured 0 to 12.8 s after the offset of SW events. For each epoch, a first-pass artifact correction procedure that identified large amplitude artifacts was performed by searching in 200 ms sliding windows for peak-to-peak voltage changes of 500 µV within each channel, and interpolating any segment that crossed this threshold using non-artifact time points before and after the segment. If more than 25% of segments of a time series of a channel was marked for correction, then the entire epoch for that channel was interpolated using data from neighboring channels. If 80% or more of the channels exceeded the 25% segment threshold, then the epoch was discarded entirely. A second-pass artifact correction was then performed such that any channel that exceeded the 500 µV (peak-to-peak voltage change) threshold across the time series within a trial was reconstructed by interpolation of its neighbors.

Following artifact correction, trials were selected with the constraint that each trial had enough usable data both pre- and post-SW event to have good time-frequency estimates of the lowest frequency of interest (i.e., 0.5 Hz); otherwise, they were rejected. All epochs were then truncated to -6.4 to -1 s before the SW event and 3 to 12.8 s after the SW event in order to avoid high-amplitude stimulation artifacts. This same truncation was applied to data from the Sham nights. Finally, all epochs were mean centered, bandpass filtered with a Butterworth filter between 0.1 and 125 Hz, and bandstop filtered between 59 and 61 Hz, and all channels were re-referenced to the global average across channels.

Time frequency decomposition of the data was performed in FieldTrip. Prior to decomposition, symmetric (mirror) padding was applied to extend the pre- and post-SW event time series to reduce edge artifacts. EEG epochs were then convolved with Morlet wavelets starting with a width of 4 at the center frequency of 0.5 Hz and increasing in width logarithmically up to a maximum width of 7 in order to minimize the combined uncertainty in time and frequency domains. Simultaneously, subsequent center frequencies were chosen such that each wavelet was one standard deviation in frequency domain from the previous wavelet. This process resulted in a time-frequency representation of roughly 35 log-spaced frequency bins from 0.5 to 50 Hz and equally-spaced time bins of 20 ms. Once time-frequency data was calculated, pre-SW event data from -3.5 to -3 s in each frequency bin was concatenated across trials and used to estimate a mean and standard deviation. These values were then used as a baseline to z-score both the pre- and post-SW event power for each trial and within each frequency bin to compute spectral power changes without single-trial bias (Ciuparu & Mureşan, 2016). These values were then averaged across trials within Active and Sham nights to yield a single channel x time x frequency matrix per condition for each subject. Subject averages were created using a random subset of trials such that trial numbers were matched between the Active and Sham nights.

Significant differences in spectral power changes and correlations with behavior were assessed statistically using non-parametric cluster-based permutation tests (Maris & Oostenveld, 2007). Averaged spectral power changes were calculated within pre-defined frequency bands: slow-wave (0.5-1.2 Hz), delta (1-4 Hz), theta (4-8 Hz), slow-spindle (9-12 Hz), and fast-spindle (12-15 Hz). For each frequency band, a paired-sample t-test was performed at each channel x time bin between 3 and 10 s from the offset of SW events, and clusters were created by grouping adjacent channels and time bins that had a *P* value < 0.05. Each cluster was then characterized by the sum of its t-values, and cluster-level statistics were evaluated using a permutation distribution created by shuffling the subject labels and repeating the clustering procedure 2000 times. Thus, a clusterwise significance value can be attributed to each observed cluster in reference to its position in the permutation-based surrogate distribution. Any cluster with a clusterwise *P* value < 0.05 after application of Bonferroni correction for 5 multiple comparisons (for five frequency band examinations) was considered significant.

Significant clusters from the first analysis (called “contrast clusters”) were then used as a mask to perform a second cluster-based permutation test on the correlation between overnight metamemory changes and previously significant channel x time bins. This effectively limited the correlation cluster analysis to the channel x time bins that showed an *a priori* significant difference between the Active and Sham nights. At each significant channel x time bin, a correlation coefficient between spectral power changes and overnight metamemory changes was calculated. The computed correlation coefficients were transformed into t-values, which were used to calculate *P* values. The same permutation-based significance test was performed to group adjacent channel x time bins with *P* values < 0.05 into so-called “correlation clusters”. Any such cluster with a clusterwise *P* value < 0.05 was considered significant.

### Supplementary Text

The metamemory scores in the five test sessions were also analyzed with a linear mixed-effects model with subject as a random factor. Fixed effects included intervention type (‘Tag & Cue’, ‘Tag & No Cue’, ‘Sham’), session (‘Test 1’, ‘Test 2’, ‘Test 3’, ‘Test 4’, ‘Test 5’), the interaction of intervention type and session, and covariates of STAMP type (‘Temporal’, ‘Spatial’), stimulation condition order (‘Active First’, ‘Sham First’), and episode group (‘Group A’, ‘Group B’). Metamemory significantly differed across sessions (F(4,336) = 3.0569, *p* = 0.01701), and there was a marginally significant interaction between intervention type and session (F(8,336) = 1.7941, *p* = 0.07721). All other effects were not significant.

We also investigated the effects of STAMPs on sensations during episode viewing using a repeated measures ANOVA with 2 within-subjects factors (sensation type: ‘itching’, ‘heat’, ‘tingling’; stimulation condition: ‘Active’, ‘Sham’) and 2 between-subjects factors (STAMP type: ‘Spatial’, ‘Temporal’; stimulation condition order: ‘Active First’, ‘Sham First’). There was an omnibus effect of stimulation condition (F(1,20) = 28.498, *p* = 0.000032), suggesting that when collapsed across sensation types, sensations during Active stimulation were rated higher by 1.581 compared to sham. All other effects were not significant, including between tDCS and tACS STAMPs. Numerically tACS STAMP sensation values were rated on average lower than tDCS STAMP sensation values when collapsed across within-subject factors. The overall mean values for sensations were all below 2.5 out of 10 in the Active stimulation condition and below 1 out of 10 in the Sham stimulation condition, suggesting STAMPs were well tolerated.

Further, we ran a series of chi-squared tests to investigate whether subjects were blind to the stimulation conditions. Due to the within-subjects nature of the design, we classified subjects by whether or not they were able to guess both stimulations conditions successfully (*χ*^2^(1) = 0.053, *p* = 0.818546). We then looked at each stimulation condition separately, regardless of order. For the Active stimulation condition, subjects were not able to guess their condition successfully (*χ*^2^(1) = 0.182, *p* = 0.669815). For the Sham stimulation condition, however, all subjects guessed their condition successfully; so a chi-squared test could not be performed. Because of this, we looked at stimulation condition order effects (‘Active First’ vs. ‘Sham First’). ‘Active First’ subjects were not to be guess their stimulation conditions successfully (*χ*^2^(1) = 1.600, *p* = 0.205903), whereas ‘Sham First’ subjects were able to do so at a trend level (*χ*^2^(1) = 2.778, *p* = 0.095581). These results suggest that the subjects were sufficiently blind to their stimulation condition and order assignments.

---

**Fig. S1.**
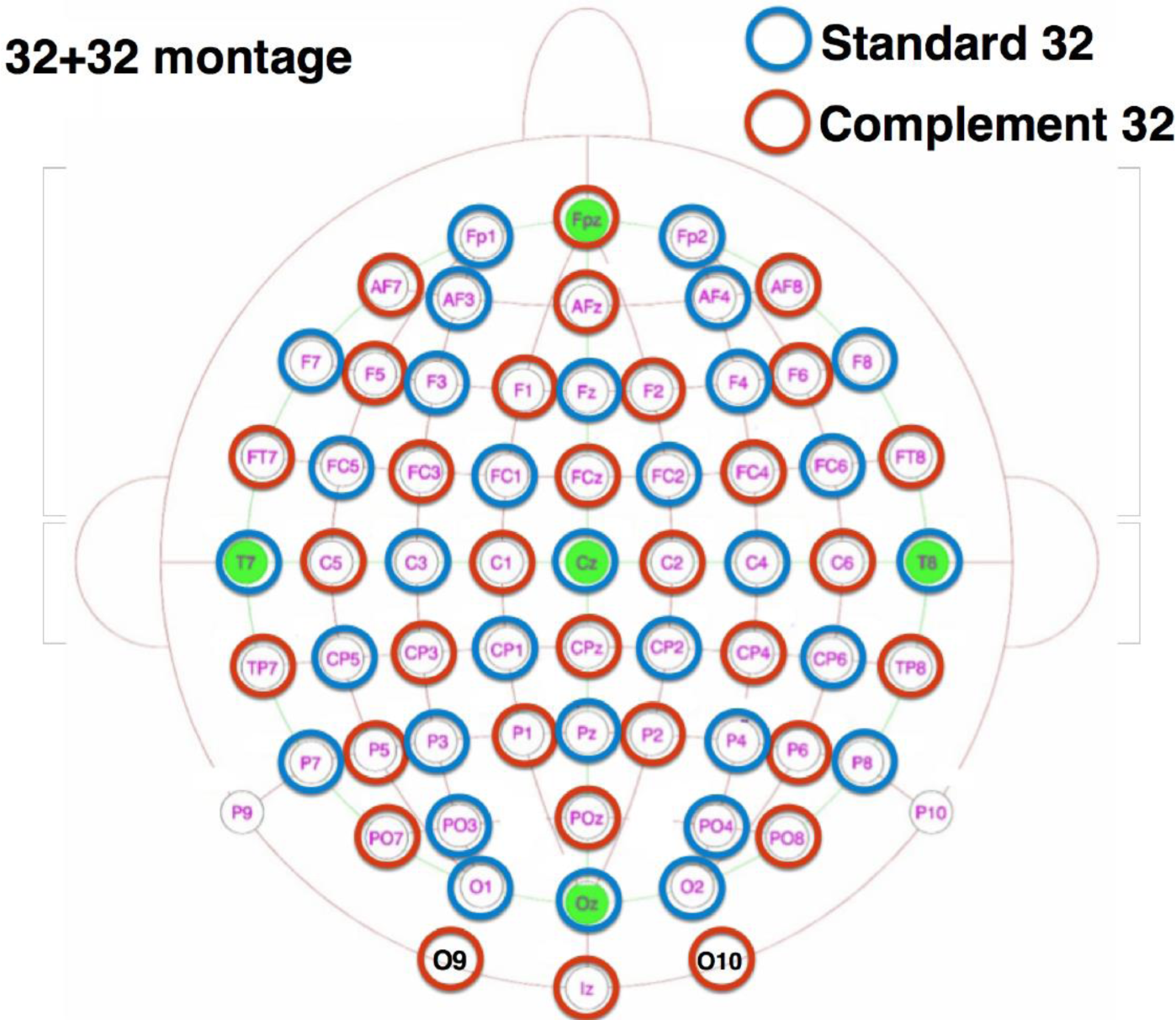
Scalp layout of the 32 EEG and 32 stimulation electrodes used in the waking and sleep interventions. The EEG electrodes were placed according to the international 10-20 system.

**Fig. S2.**
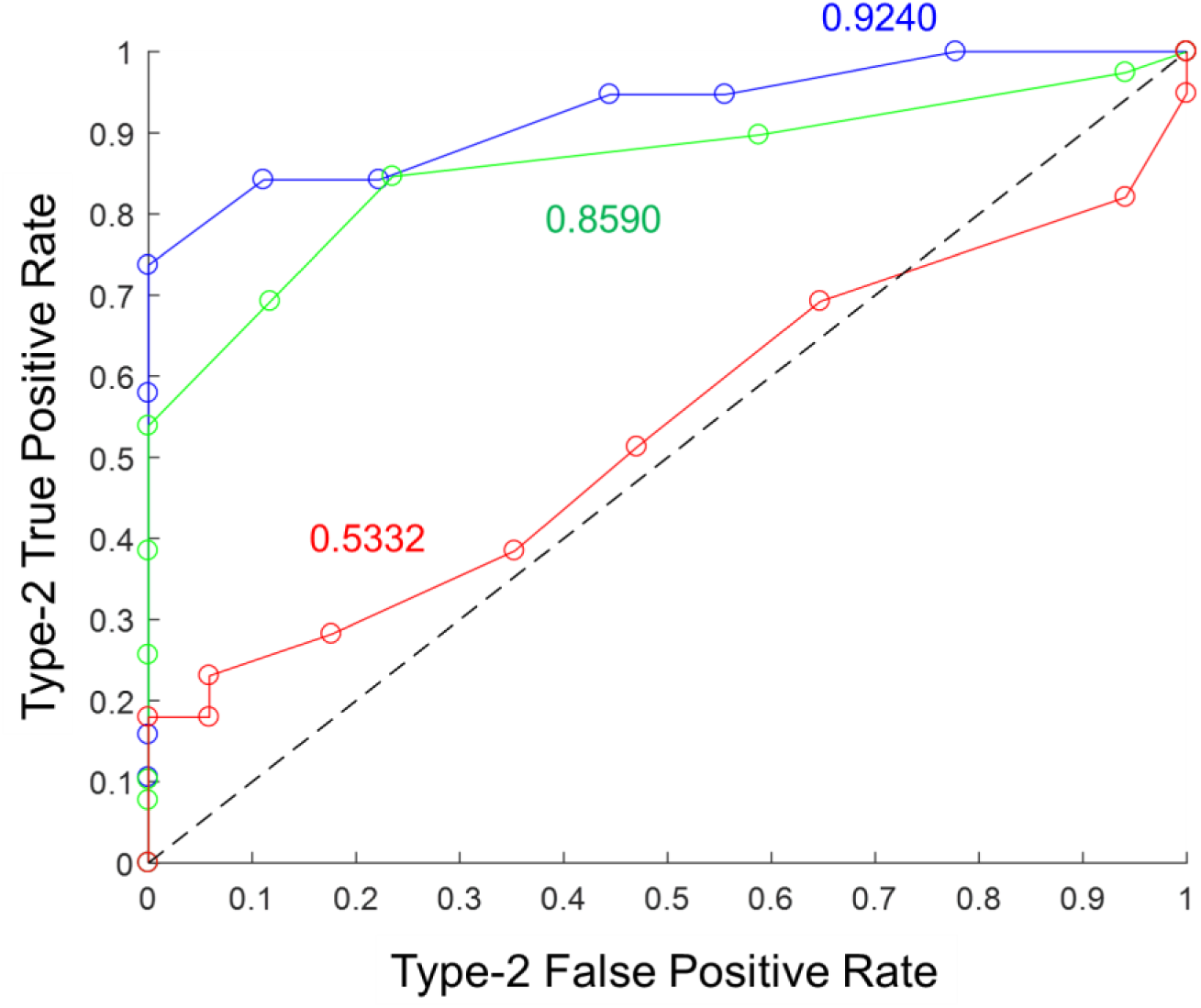
Illustration of Type-2 ROC analysis to assess episodic metamemory. Type-2 ROC curves from the three days of testing are shown for a representative subject in the Sham stimulation condition. Blue corresponds to Day 1, green to Day 2, and red to Day 3. The corresponding metamemory score (calculated as the area under the curve) is shown in the matching color.

**Table S1.** Names of the episodes in each of the four subgroups that were used in the experiment.

| **Episode Group A** | **Episode Group B** |
| --- | --- |
| Bomb Drop | Call the Car |
| Car Bomb Assassination | Car Bomb Attack |
| Cardiac Arrest | Domestic Incident |
| Deadly Argument | Drive By |
| Failed Rescue | Exposed Rendezvous |
| Handoff | Fire Response |
| Helicopter Scare | Missile Strike |
| Interrupted Meeting | Shopping |
| Missed the Bus | Sniper |
| No Escape | Street Hawker |
| Pick Pocket | Suicide Bomb |
| Repair Man | The Dare |
| Timely Exit | The Meeting |
| Waiting Gunman | The Serenade |

## Movie captions

**Movie S1.**

Video of the “Fire Response” episode used in the experiment.

**Movie S2.**

Video demonstration of the GUI that was used to administer the quizzes for assessing memory performance.

## Supplementary Note

This section provides details for each of the 28 episodes that were used in the experiment.

(1) Bomb Drop

Textual prompt: In the episode where two characters meet in front of the building, one enters the building, and an explosion occurs

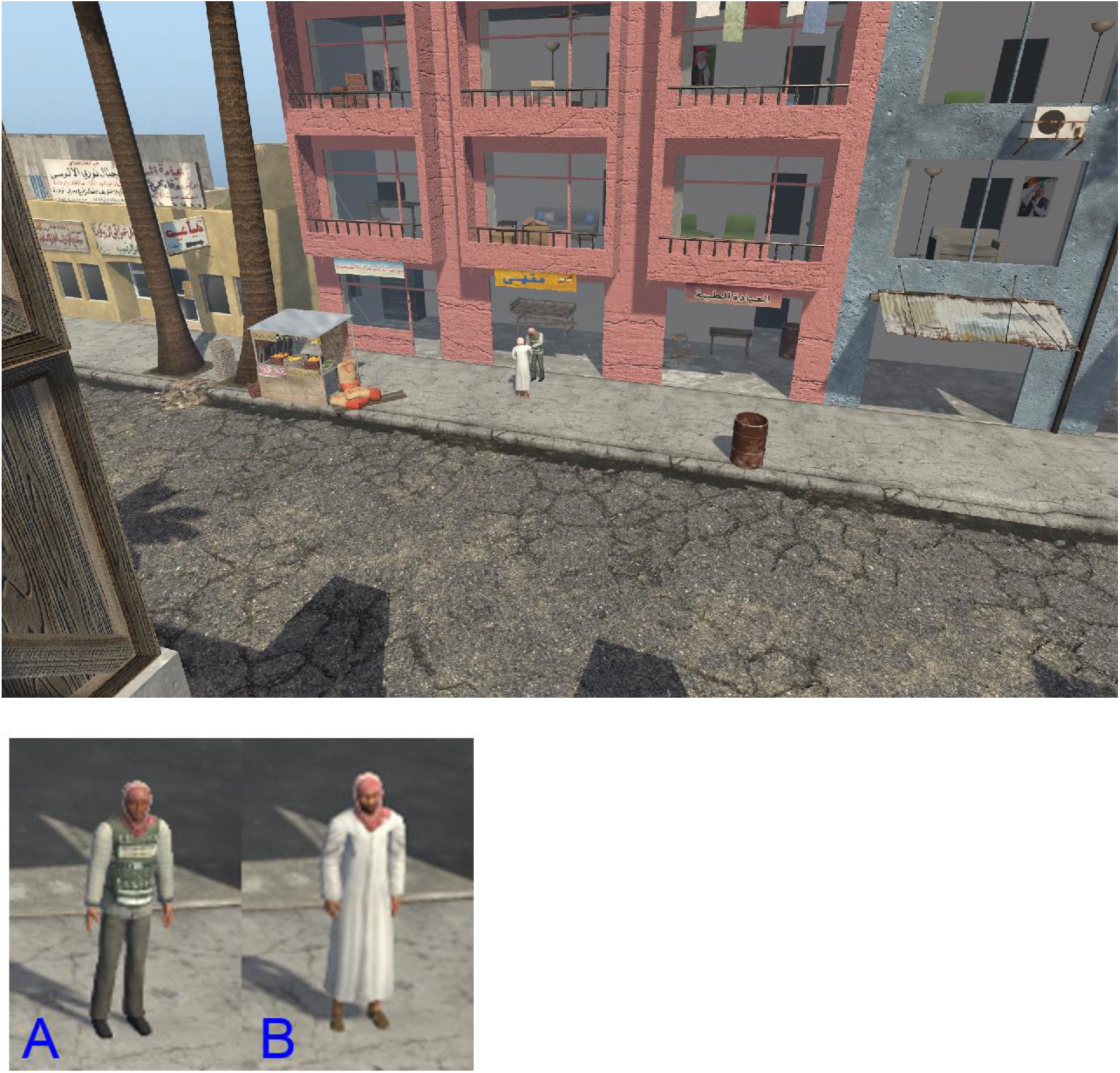

Question 1: The bomb exploded on the 2nd floor

Answer: False

Question 2: Character A gets out of a car

Answer: False

Question 3: Character A exits the building before an explosion occurs

Answer: True

Question 4: Character A fell down when the explosion occurred

Answer: False

Question 5: Character B runs away from the building (to the right) before the blast

Answer: True

Question 6: Character A was inside building when blast occurred

Answer: False

Question 7: Character B enters car and drives away

Answer: False

Question 8: Character A enters the building after interacting with Character B on the sidewalk

Answer: True

Question 9: Character B runs away from building before explosion occurs

Answer: True

Question 10: Neither Character A nor Character B came from the building

Answer: True

(2) Call The Car

Textual prompt: In the episode where one character arrives in a vehicle, converses with the other, then they both leave in the vehicle

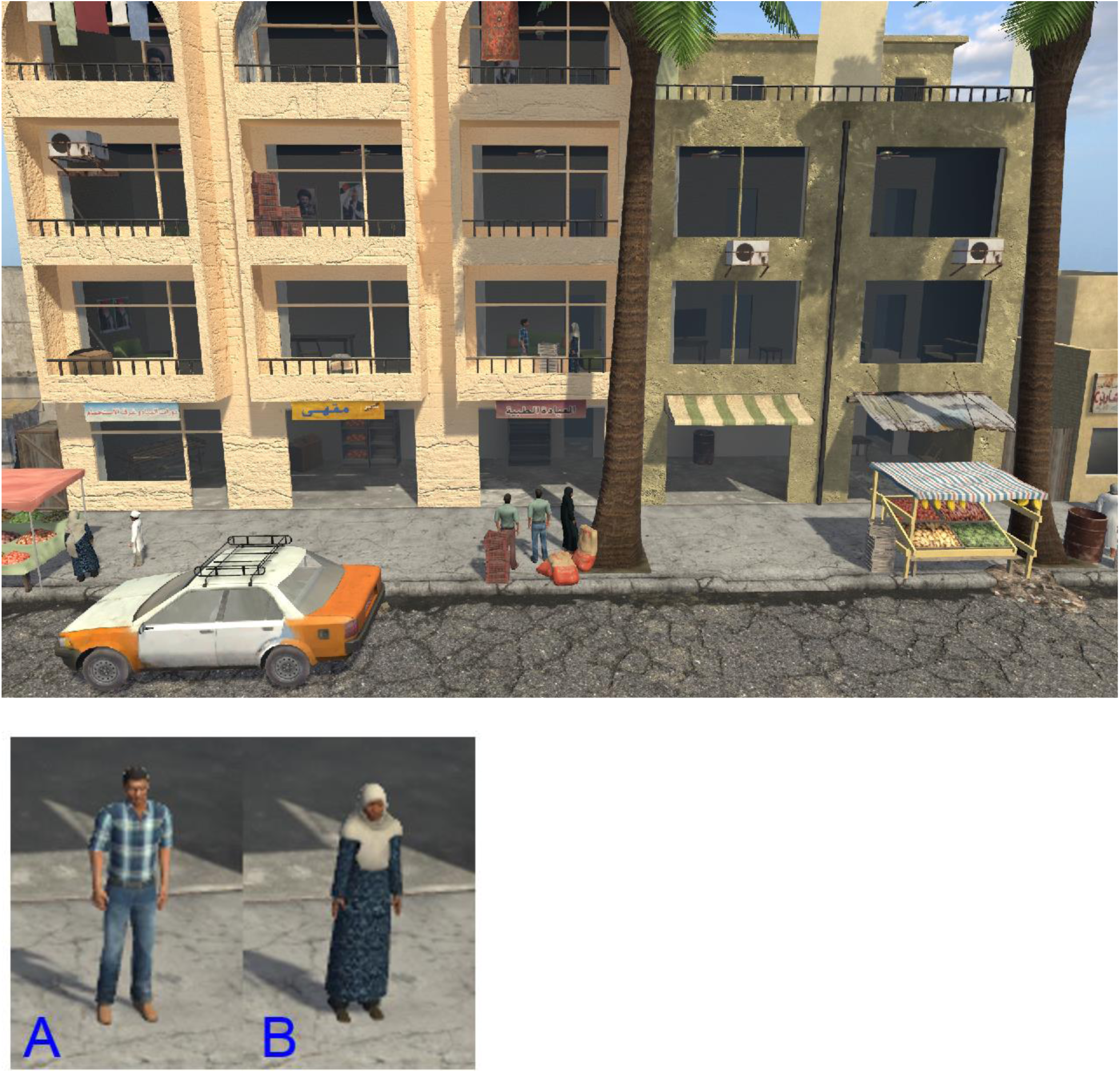

Question 1: Character B looked out a window before a car arrived

Answer: True

Question 2: Character A got out of a car and ran into the building

Answer: True

Question 3: The two characters met on the street

Answer: False

Question 4: Character B is only seen in a single room before being met by character A

Answer: True

Question 5: CharatersA and B converse a second time on the sidewalk

Answer: False

Question 6: Characters A and B converse in multiple rooms in the building

Answer: False

Question 7: Characters A and B left the building through the same exit

Answer: True

Question 8: A police car arrived

Answer: False

Question 9: A helicopter flew over while Characters A and B were conversing

Answer: False

Question 10: Character B followed Character A to the car

Answer: True

(3) Car Bomb Assassination

Textual prompt: In the episode where several characters converse inside the building, exit the building, and the police car explodes after some get into the vehicle

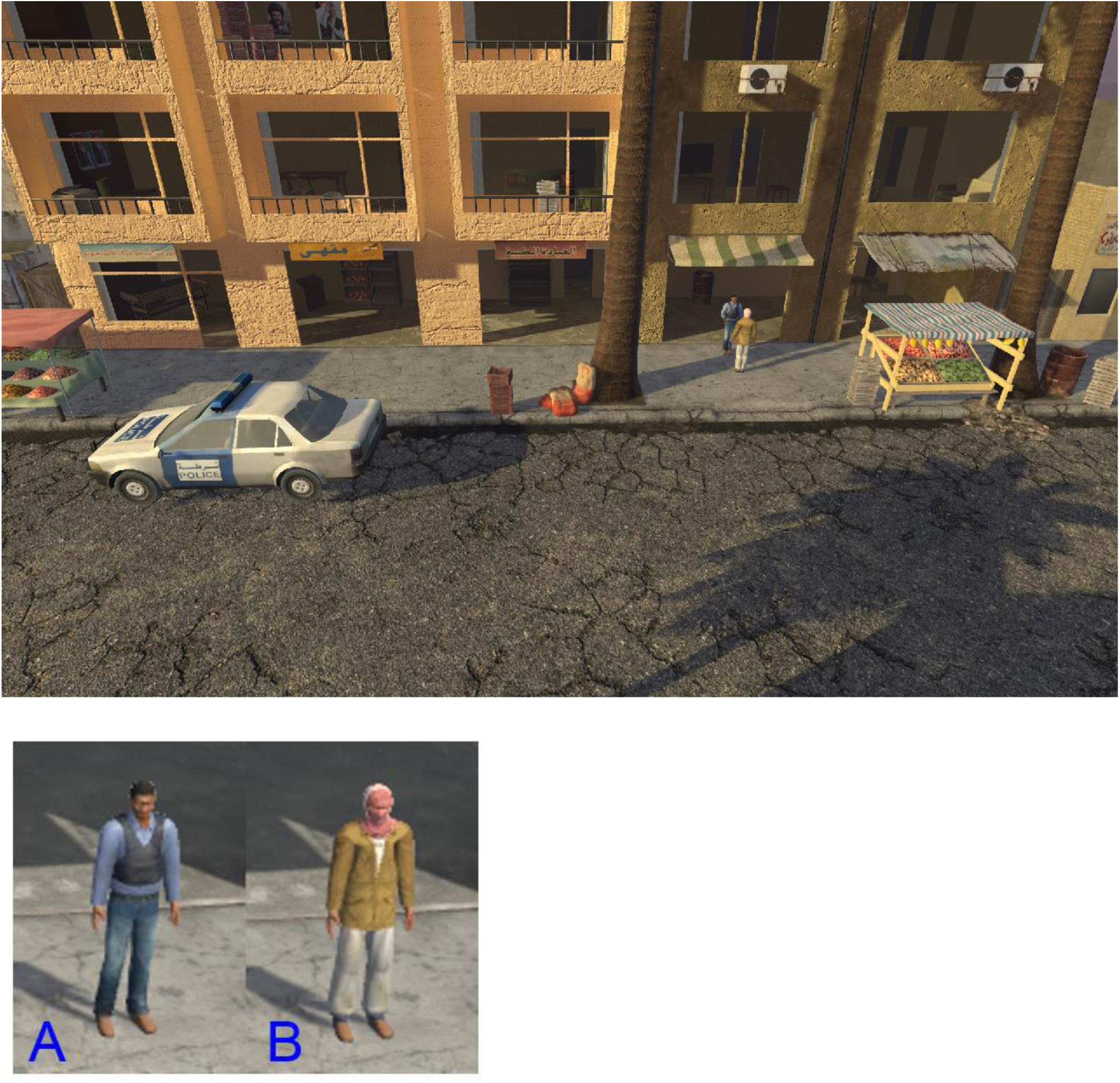

Question 1: The characters met inside the building

Answer: True

Question 2: The characters left the building and went in different directions

Answer: False

Question 3: A person was seen running from the car before it exploded

Answer: False

Question 4: The car burned after it blew up

Answer: True

Question 5: There was only one vehicle in the scene

Answer: False

Question 6: An ambulance pulled up after the vehicle exploded

Answer: True

Question 7: Character A and B exit the building together

Answer: True

Question 8: The car started smoking shortly before exploding

Answer: False

Question 9: Character A runs down the street after the explosion

Answer: False

Question 10: Gunshots are heard before explosion

Answer: False

(4) Car Bomb Attack

Textual prompt: In the episode where a car arrives, one of the characters exits the car, and the car explodes

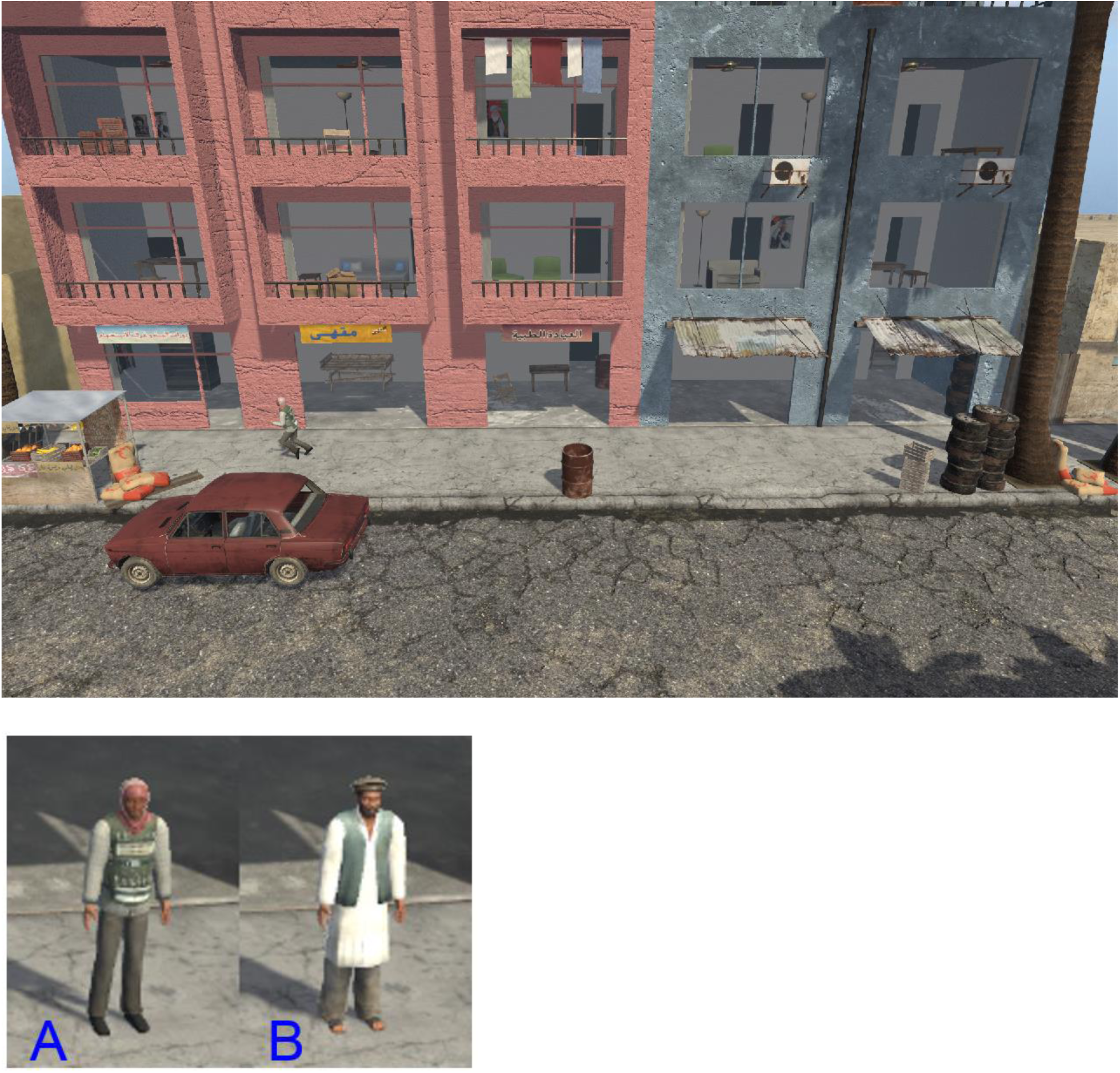

Question 1: Character B entered the building just before a car arrived

Answer: True

Question 2: Character A ran out of the car and into the building

Answer: False

Question 3: Character B left the building before the car blew up

Answer: False

Question 4: Characters A and B conversed before the explosion

Answer: False

Question 5: Character B ran away from the building after the explosion

Answer: True

Question 6: Character A falls down on street when explosion occurs

Answer: False

Question 7: Character A arrives in a police vehicle

Answer: False

Question 8: Fire starts in the building

Answer: True

Question 9: Character A does not run into the building before the car explodes

Answer: True

Question 10: The car catches fire after the explosion

Answer: True

(5) Cardiac Arrest

Textual prompt: In the episode where one character falls over on the sidewalk, another goes inside, and an ambulance arrives

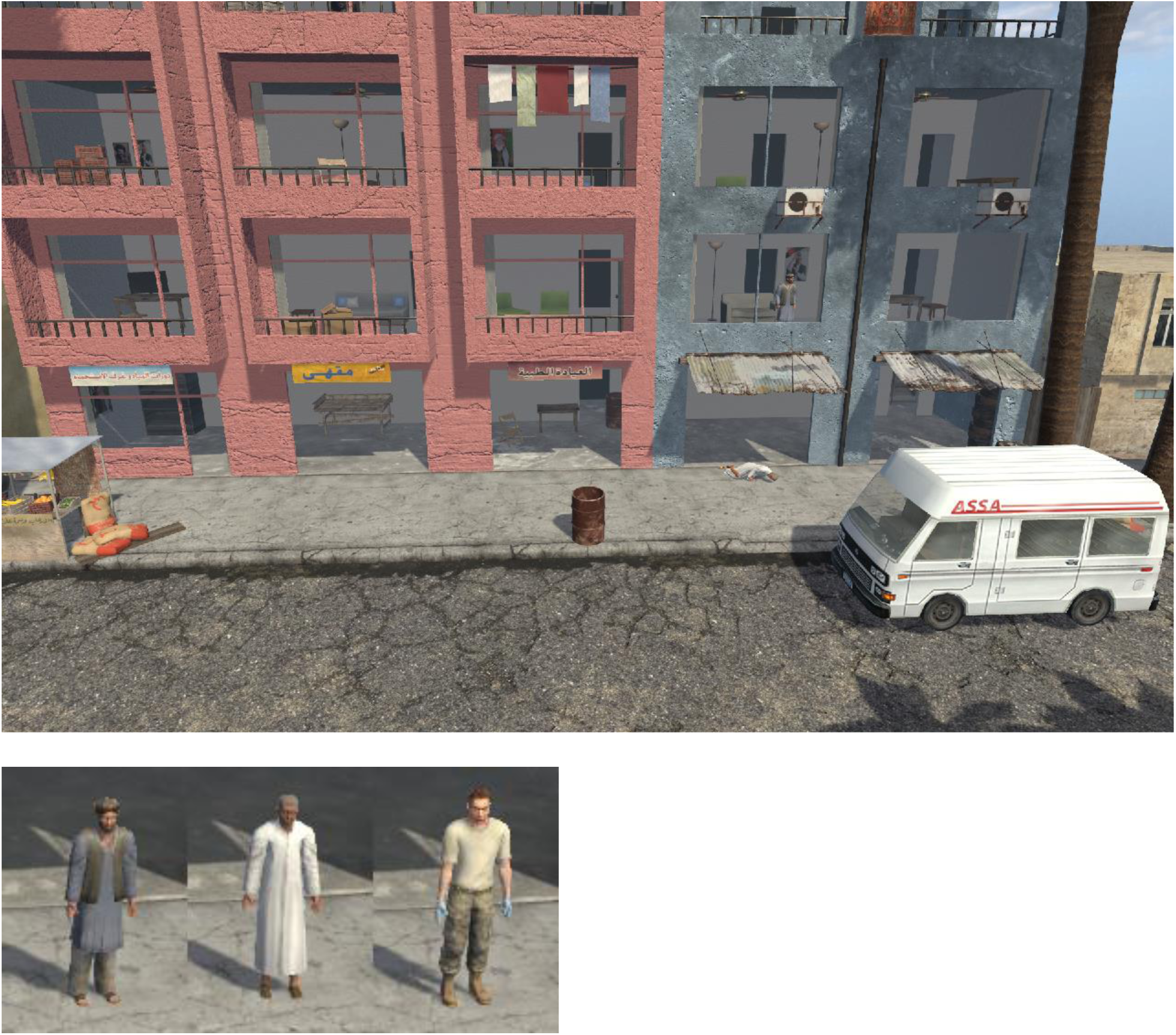

Question 1: Character B is loaded into the ambulance

Answer: False

Question 2: Character B stands back up after the ambulance arrives

Answer: False

Question 3: Gunshots are heard before Character B falls down

Answer: False

Question 4: Character C arrives before the ambulance

Answer: False

Question 5: Characters A and B exit the building start talking on the sidewalk

Answer: True

Question 6: Character A does not come back downstairs after the ambulance arrives

Answer: True

Question 7: A police car arrives

Answer: False

Question 8: Character C gets out of an ambulance to help Character B

Answer: True

Question 9: After the ambulance arrives, Character A leaves the building and runs down the sidewalk

Answer: False

Question 10: Character A closes the blinds after the ambulance arrives

Answer: False

(6) Deadly Argument

Textual prompt: In the episode where two characters have a disagreement and one shoots the other in the building

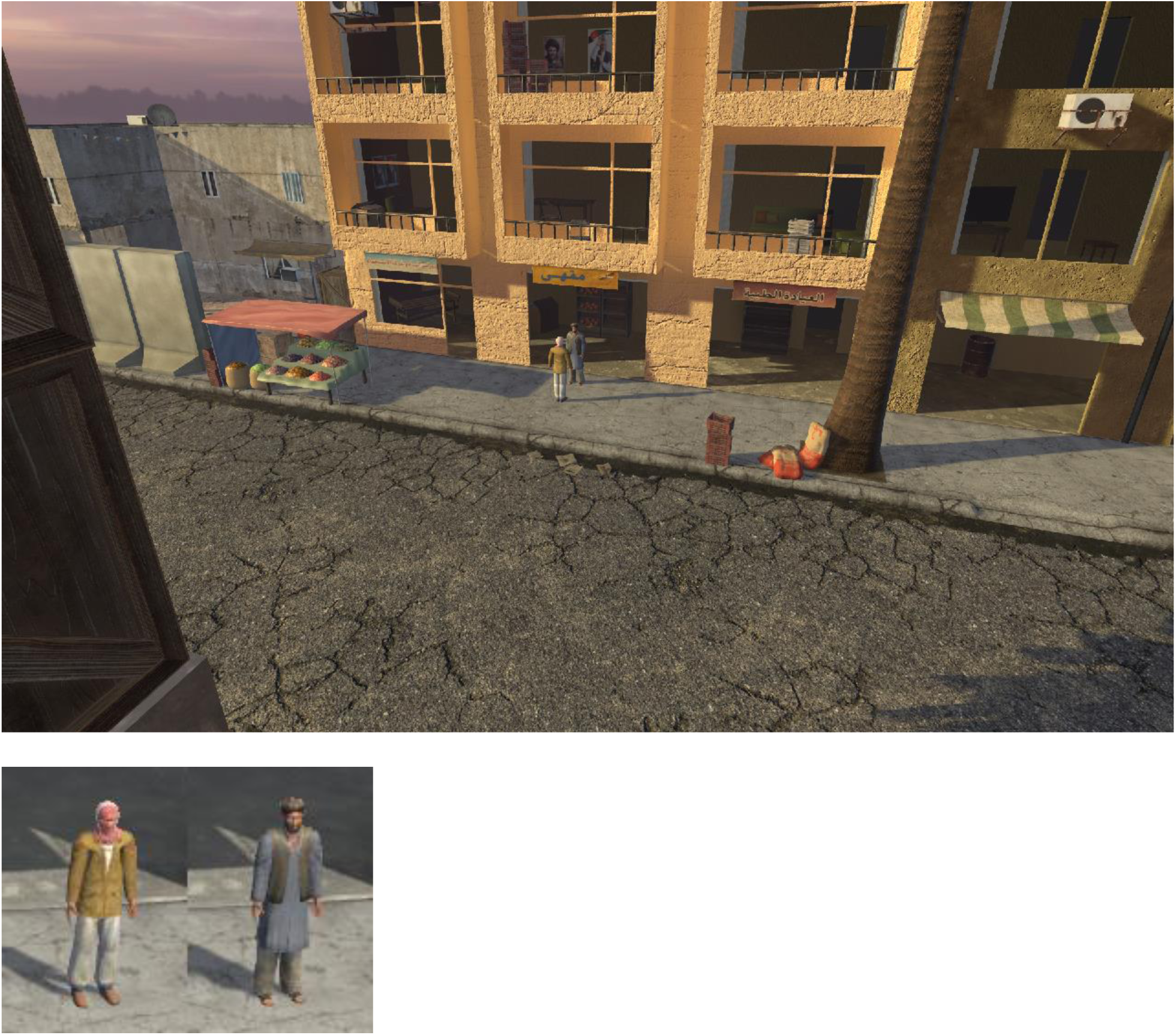

Question 1: Characters A and B walked into view together

Answer: False

Question 2: Character B entered the building while Character A walked away from him

Answer: True

Question 3: Character A enters the building with Character B

Answer: False

Question 4: Characters A and B converse again inside the building

Answer: False

Question 5: Character A ran from the building after shooting Character B

Answer: True

Question 6: Characters A and B entered the building together

Answer: True

Question 7: After Character B fell to the floor, Character A brought him outside the building

Answer: False

Question 8: Characters A and B were seen together in a room in the middle of the building

Answer: True

Question 9: Character A shot Character B from a different room

Answer: False

Question 10: An ambulance drove up to the front of the building

Answer: False

(7) Domestic Incident

Textual prompt: In the episode where two characters have a disagreement and one shoots the other inside the building

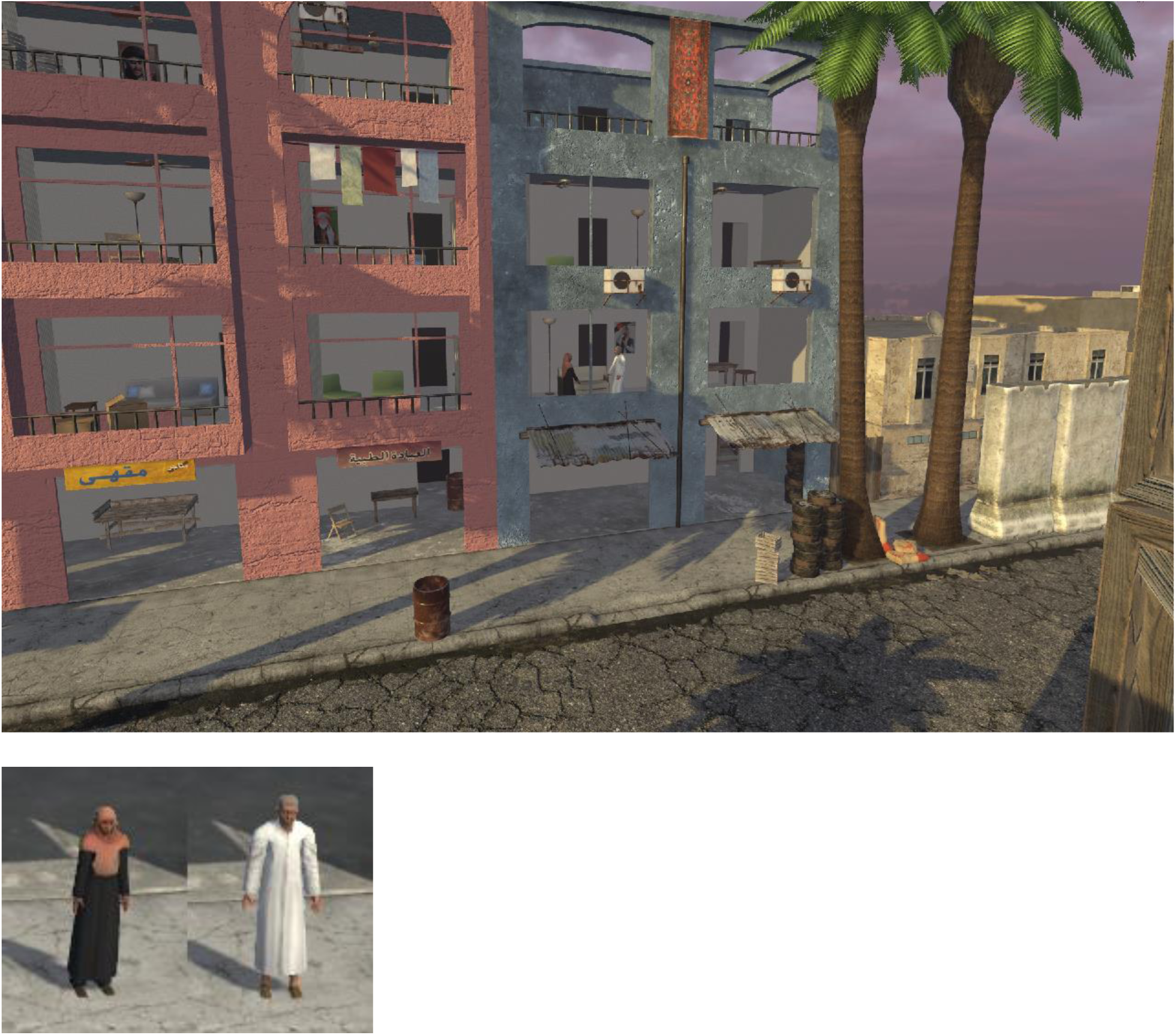

Question 1: Character A stays in the room she is in while Character B goes into another room

Answer: True

Question 2: Character B speaks to Character A in a room on the second floor

Answer: True

Question 3: A police car pulled up while Characters A and B were in separate rooms

Answer: False

Question 4: Character B left the room and then came back

Answer: True

Question 5: Character A shot Character B

Answer: False

Question 6: Character B runs from the building after the shooting

Answer: True

Question 7: Character B fires many gun shots

Answer: False

Question 8: Character A runs down the street away from the building

Answer: False

Question 9: Character B arrives by car

Answer: False

Question 10: Character B shoots Character A from behind

Answer: True

(8) Drive By

Textual prompt: In the episode where two characters are in front of the building as a vehicle drives past and gunshots are heard

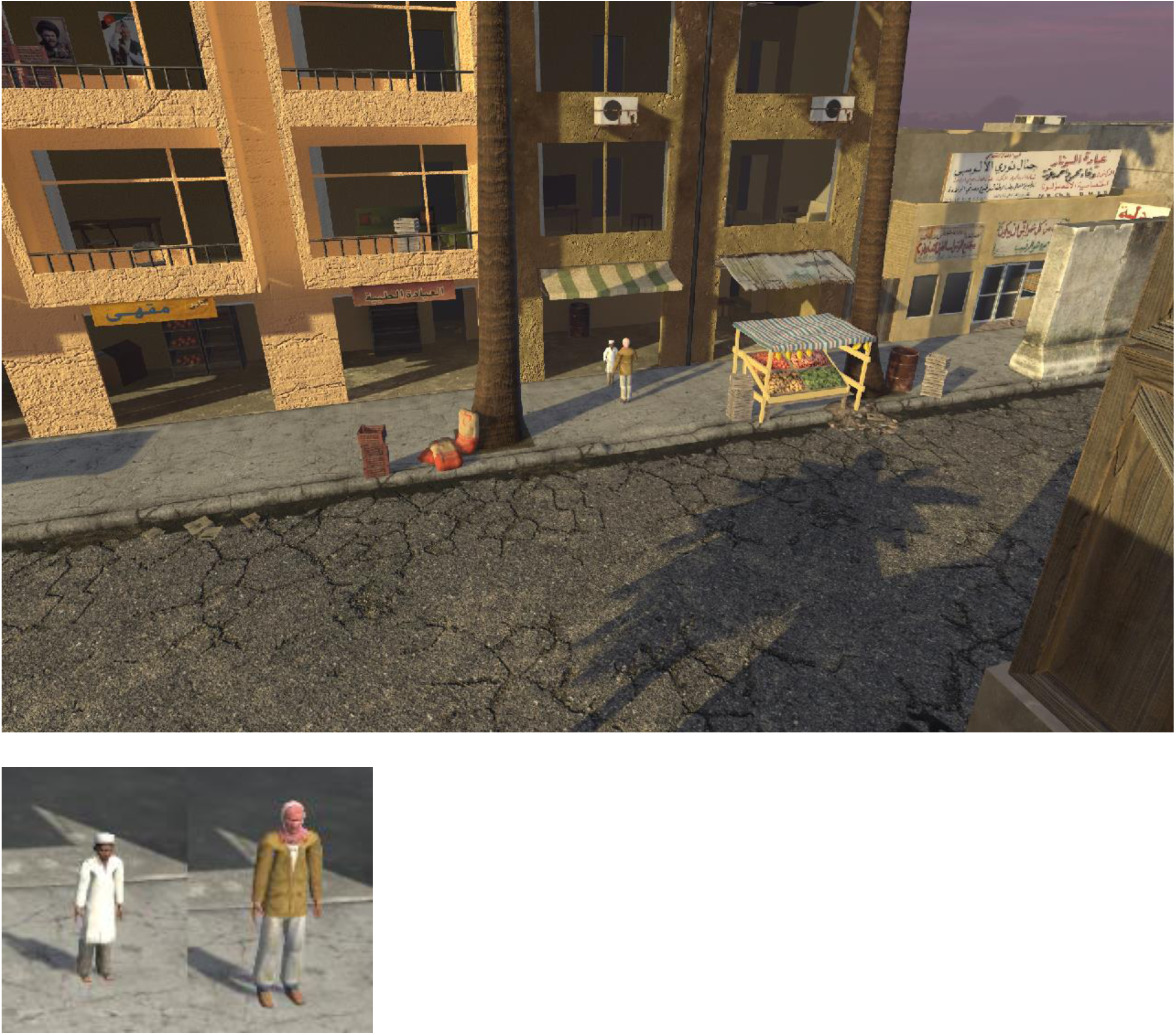

Question 1: Characters A and B were walking before stopping in front of the building

Answer: True

Question 2: A helicopter flew overhead

Answer: False

Question 3: A vehicle drove by and Character B fell to the sidewalk

Answer: True

Question 4: Both characters ran away after the vehicle passed by

Answer: False

Question 5: Only one of the people on the sidewalk fell when the truck passed

Answer: True

Question 6: Characters A and B came from the same direction

Answer: True

Question 7: Only one gunshot is heard

Answer: False

Question 8: Multiple police vehicles passed by after the first vehicle

Answer: True

Question 9: Characters A and B arrive in a car

Answer: False

Question 10: After Character A enters the building a room catches fire

Answer: False

(9) Exposed Rendezvous

Textual prompt: In the episode where two characters meet behind closed blinds, and when the blinds suddenly open, they are exposed and run away

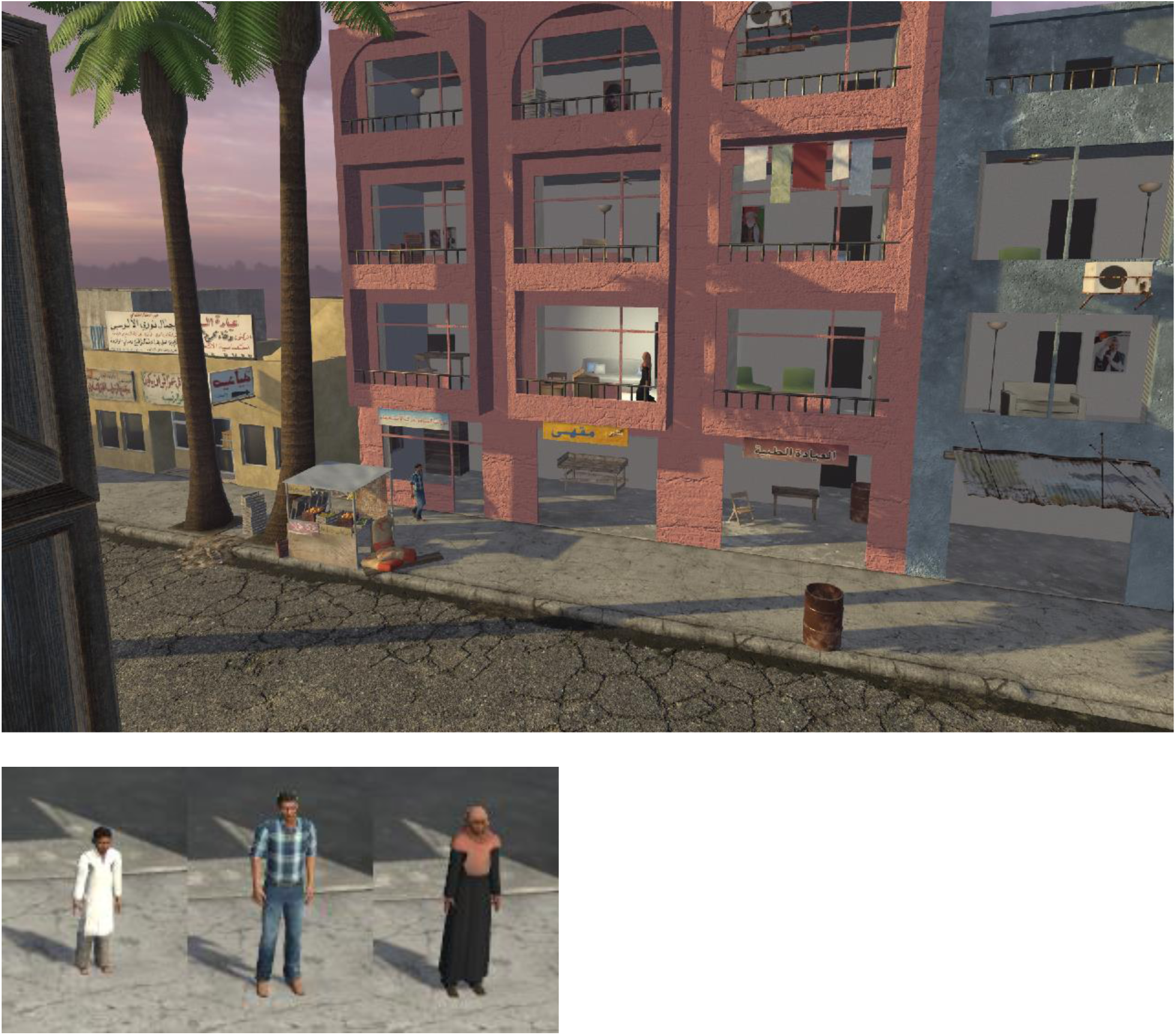

Question 1: Character C is seen entering the building

Answer: False

Question 2: Character B entered the building and is next seen moving up across rooms on every floor

Answer: True

Question 3: The curtains closed when Characters B and C are in a room together

Answer: True

Question 4: Character A is seen moving toward the room just before the curtains open again

Answer: True

Question 5: Character C looked out the window just before the curtain closes

Answer: False

Question 6: When the blinds pull back Character B and C are seen in the room

Answer: True

Question 7: Character C turns the light off when Character B is seen on the street

Answer: False

Question 8: Character A is seen in the room with Characters B and C

Answer: False

Question 9: Characters B and C leave the building and run down the street together

Answer: False

Question 10: When the curtain opens Characters B and C both look out the window

Answer: True

(10) Failed Rescue

Textual prompt: In the episode where a fire breaks out in the building while two characters are talking outside

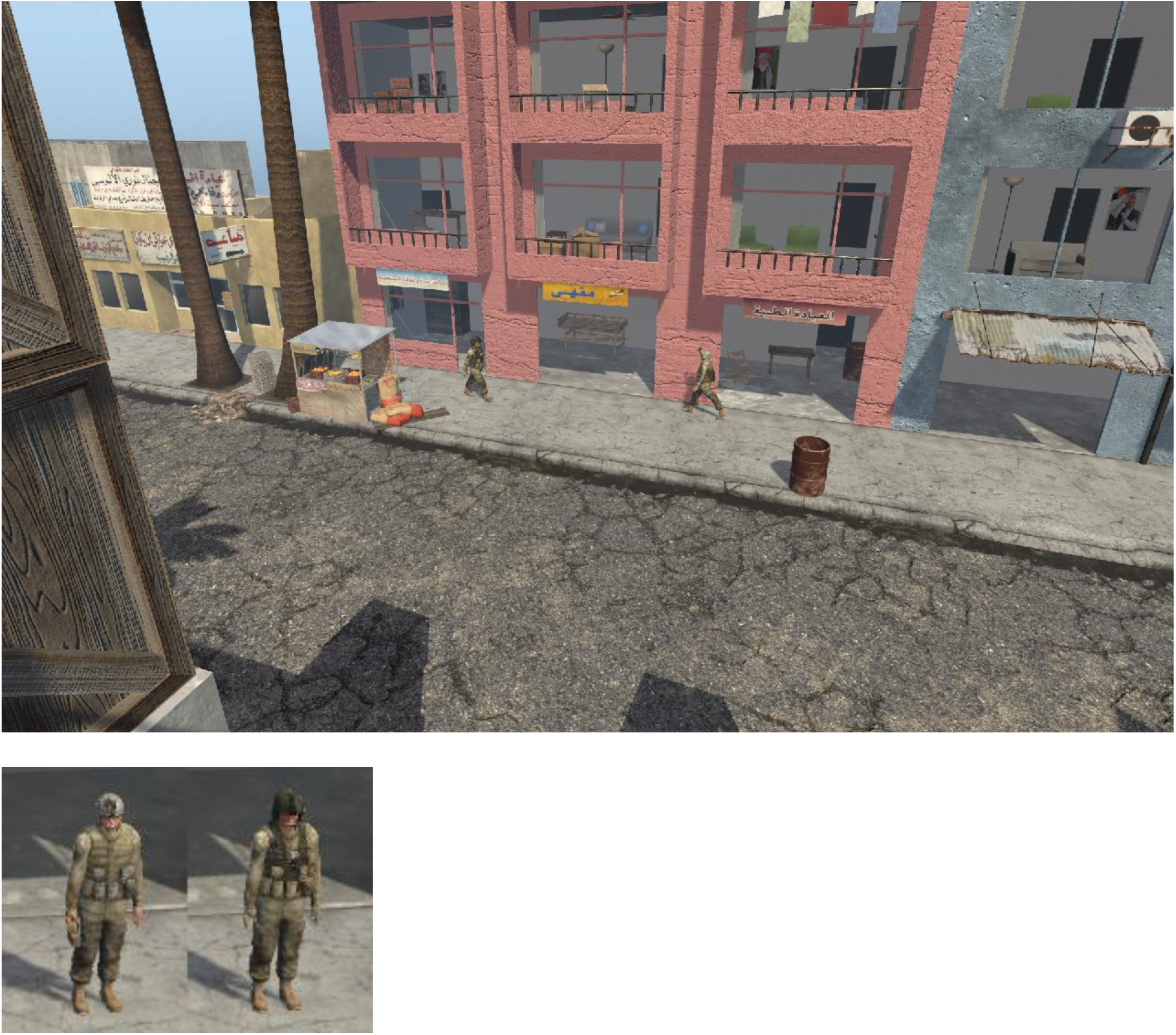

Question 1: The character that enters the building falls while the room he is in is on fire

Answer: True

Question 2: Characters A and B entered the building after fire was seen.

Answer: False

Question 3: A woman was seen on inside the building

Answer: True

Question 4: The character in the building fell down as the fire spread

Answer: True

Question 5: The fire spread to many rooms in the building

Answer: True

Question 6: Characters A and B converse in a room inside the building

Answer: False

Question 7: One soldier waited outside while the other soldier entered the building

Answer: True

Question 8: The man who did not enter the building ran away

Answer: False

Question 9: An ambulance drove up to the building

Answer: False

Question 10: A helicopter flew by during the fire

Answer: False

(11) Fire Response

Textual prompt: In the episode where two characters are in or around the building as a fire breaks out in one of the rooms and an ambulance arrives

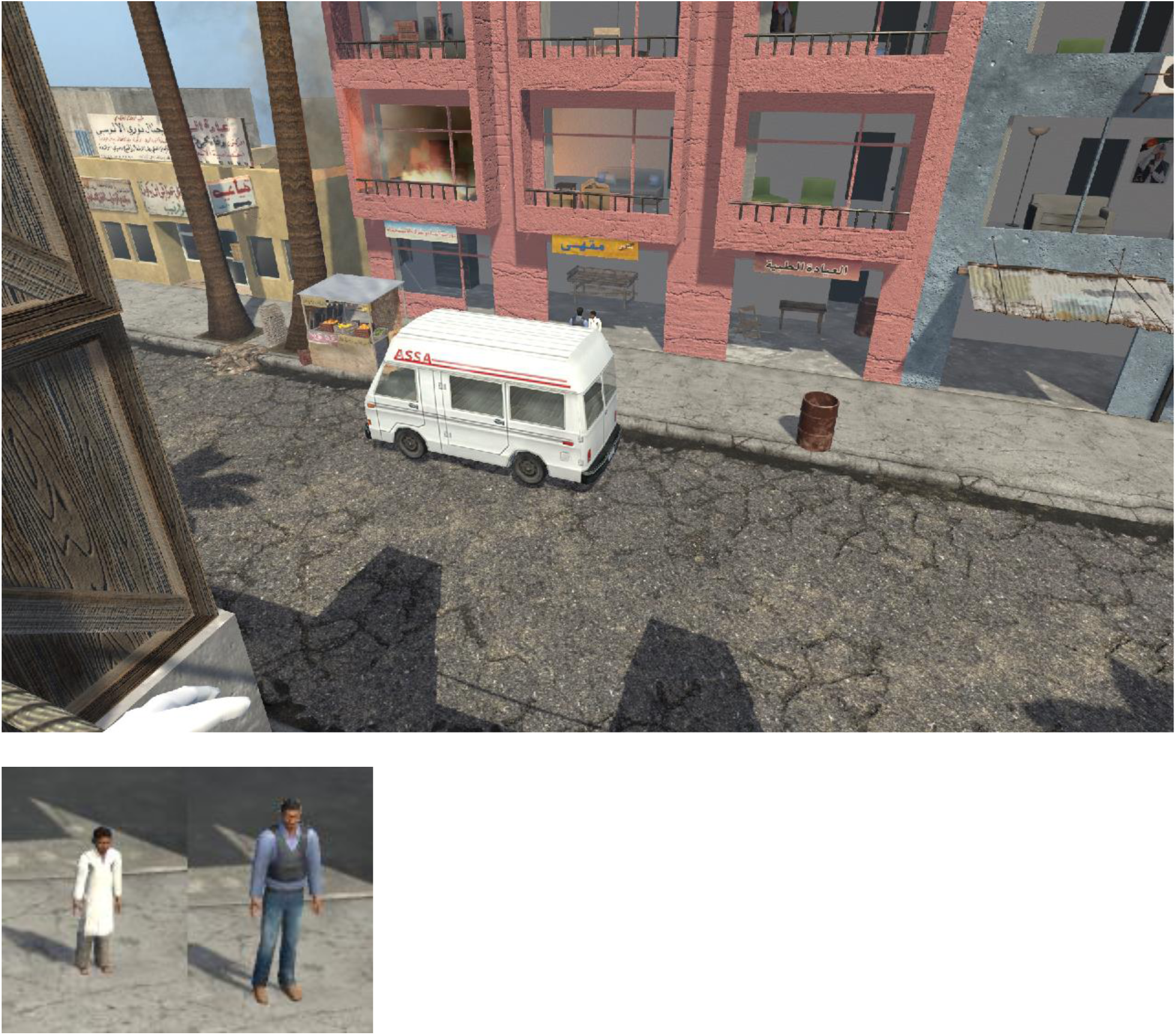

Question 1: Character B is inside the building before Character A

Answer: False

Question 2: Character B looked up at the building before entering

Answer: True

Question 3: Character A was seen in the same room as the fire

Answer: True

Question 4: Character B left the building shortly after the fire started

Answer: False

Question 5: Character B entered the room that was on fire, and left with the child in white

Answer: True

Question 6: Characters A and B talk inside the building

Answer: False

Question 7: An ambulance arrives after Characters A and B exit the building

Answer: True

Question 8: Character A got into the ambulance and it drove away

Answer: False

Question 9: Character A is inside the building before the fire starts

Answer: True

Question 10: Character A runs away after talking to Character B

Answer: False

(12) Handoff

Textual prompt: In the episode where one character is waiting in a room when the other arrives, they meet, they part, and then close the blinds

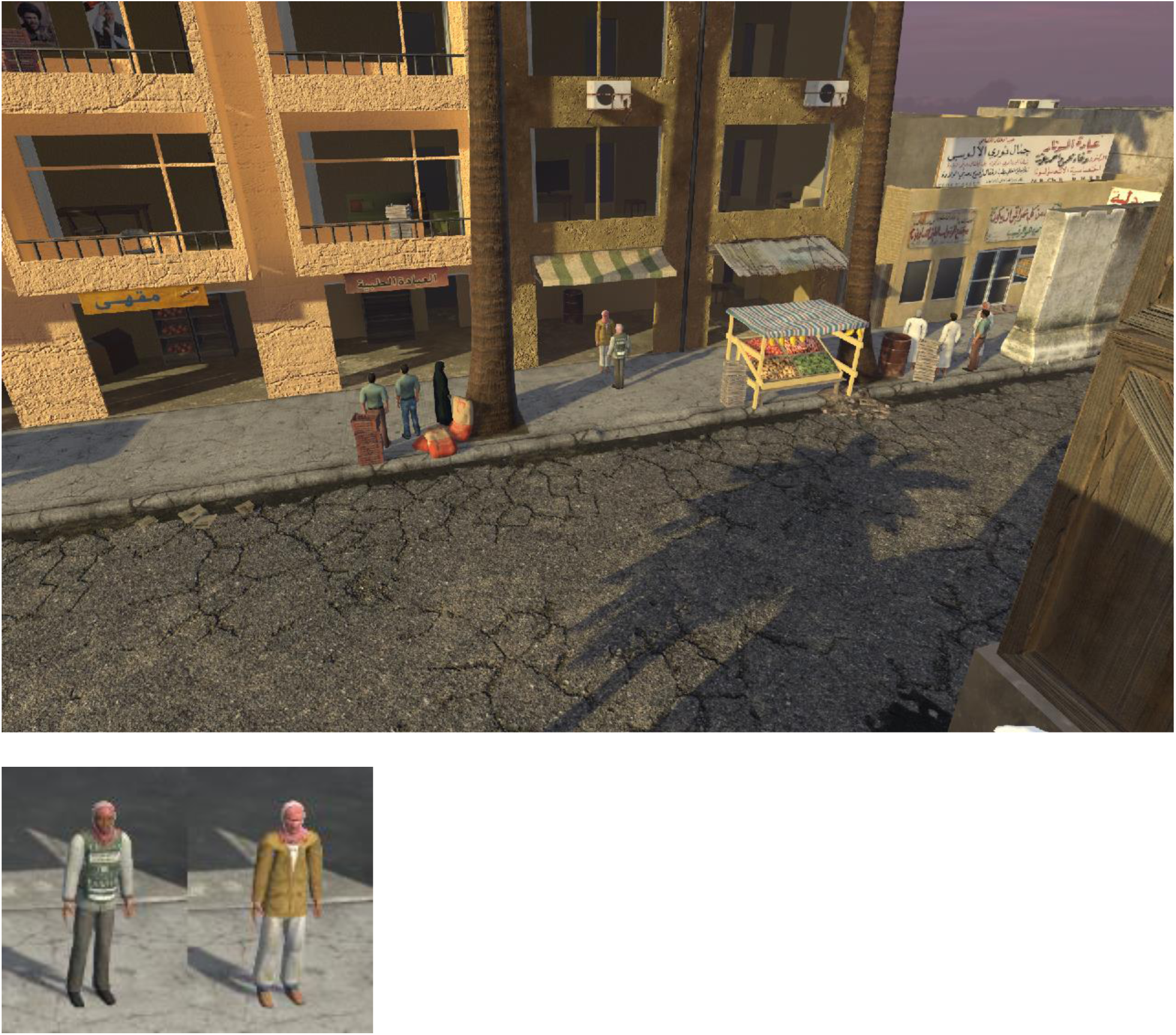

Question 1: Character B is seen pacing and then looking out from the 2nd floor

Answer: True

Question 2: Characters A and B converse inside the building before moving outside

Answer: False

Question 3: Character A leaves in a vehicle that pulls up after conversing with Character B

Answer: True

Question 4: Characters A and B meet only one enters the building

Answer: True

Question 5: When Character B gets back to the room, a police car arrives

Answer: True

Question 6: Character A enters the police car

Answer: False

Question 7: After talking to Character A, Character B walks down the sidewalk

Answer: False

Question 8: When the police car arrives, Character B goes down to meet it

Answer: False

Question 9: When Character B sees Character A on the sidewalk, Character B goes down to meet him

Answer: True

Question 10: The police car chases Character A down the street

Answer: False

(13) Helicopter Scare

Textual prompt: In the episode where a character shoots at a passing helicopter

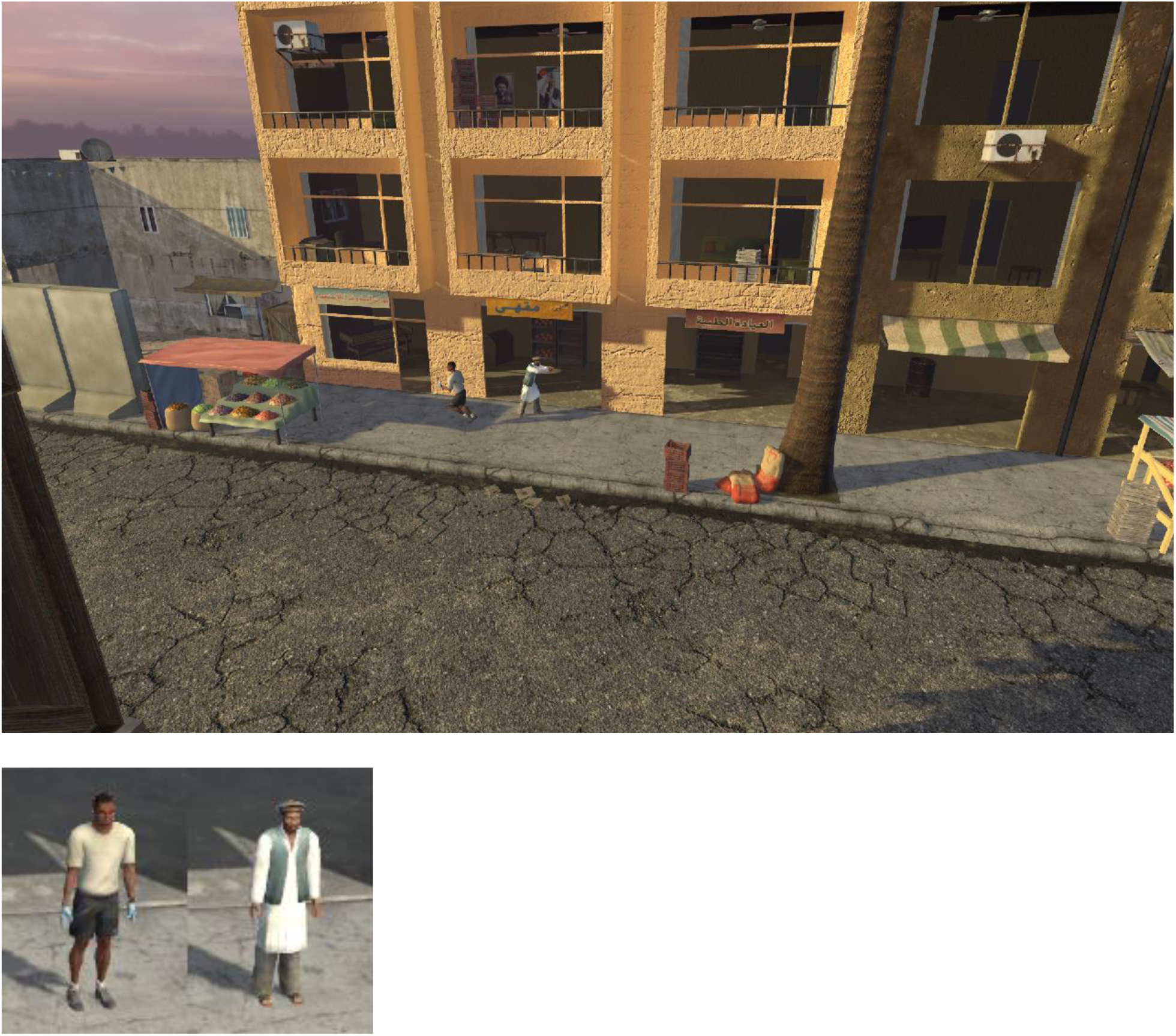

Question 1: When a helicopter is heard Character A runs inside

Answer: True

Question 2: The second time the helicopter is heard Character B runs down the sidewalk

Answer: True

Question 3: Character A ran into the left side entrance of the building

Answer: True

Question 4: An explosion occurs down the street from the building

Answer: False

Question 5: Character A exits the building after the helicopter passes by

Answer: False

Question 6: Character B takes a shooting stance when the helicopter passes overhead the first time

Answer: True

Question 7: Characters A and B ran into the building together

Answer: False

Question 8: Character B ran down the sidewalk to the right

Answer: True

Question 9: When the helicopter passes the second time an explosion occurs

Answer: True

Question 10: An ambulance drives up

Answer: False

(14) Interupted Meeting

Textual prompt: In the episode where two characters are conversing in the building, another arrives and one of the characters leaves with the new arrival

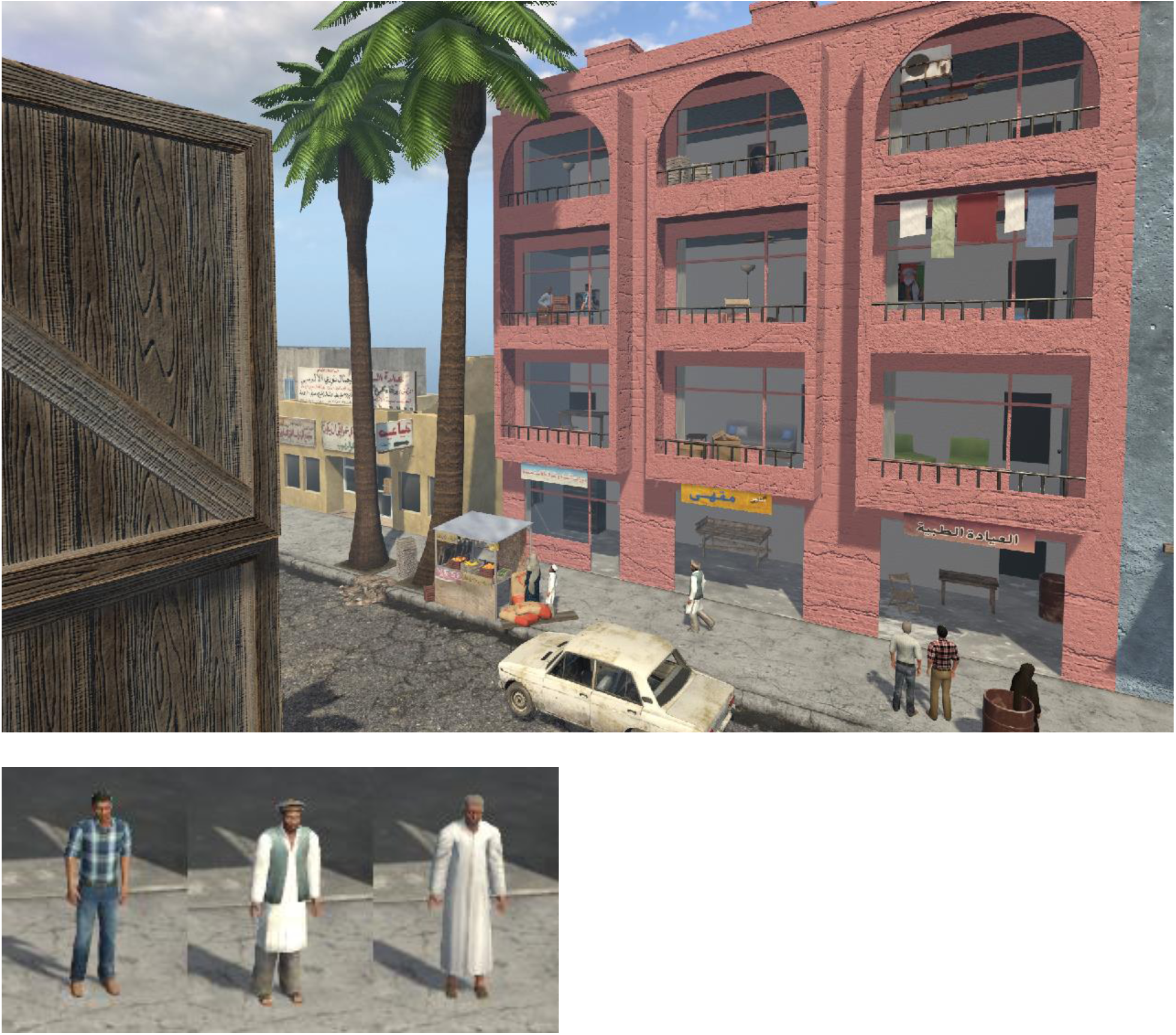

Question 1: Characters A and B are talking together in the room before Character C arrives

Answer: False

Question 2: Character B waits on the street and waits for Character A to come down

Answer: False

Question 3: A bomb explodes on the street

Answer: False

Question 4: Character B leaves the room with Character A

Answer: True

Question 5: Character C leaves and runs down the street

Answer: False

Question 6: After the two men leave in the car, the blinds close in the room

Answer: True

Question 7: Character C appears in the next room after the other two men leave

Answer: False

Question 8: Character C left the building

Answer: False

Question 9: Characters A and B converse in multiple rooms in the building

Answer: False

Question 10: Character C remains in the same room after Characters A and B depart

Answer: True

(15) Missed The Bus

Textual prompt: In the episode where two characters get into a vehicle and another runs after it down the sidewalk

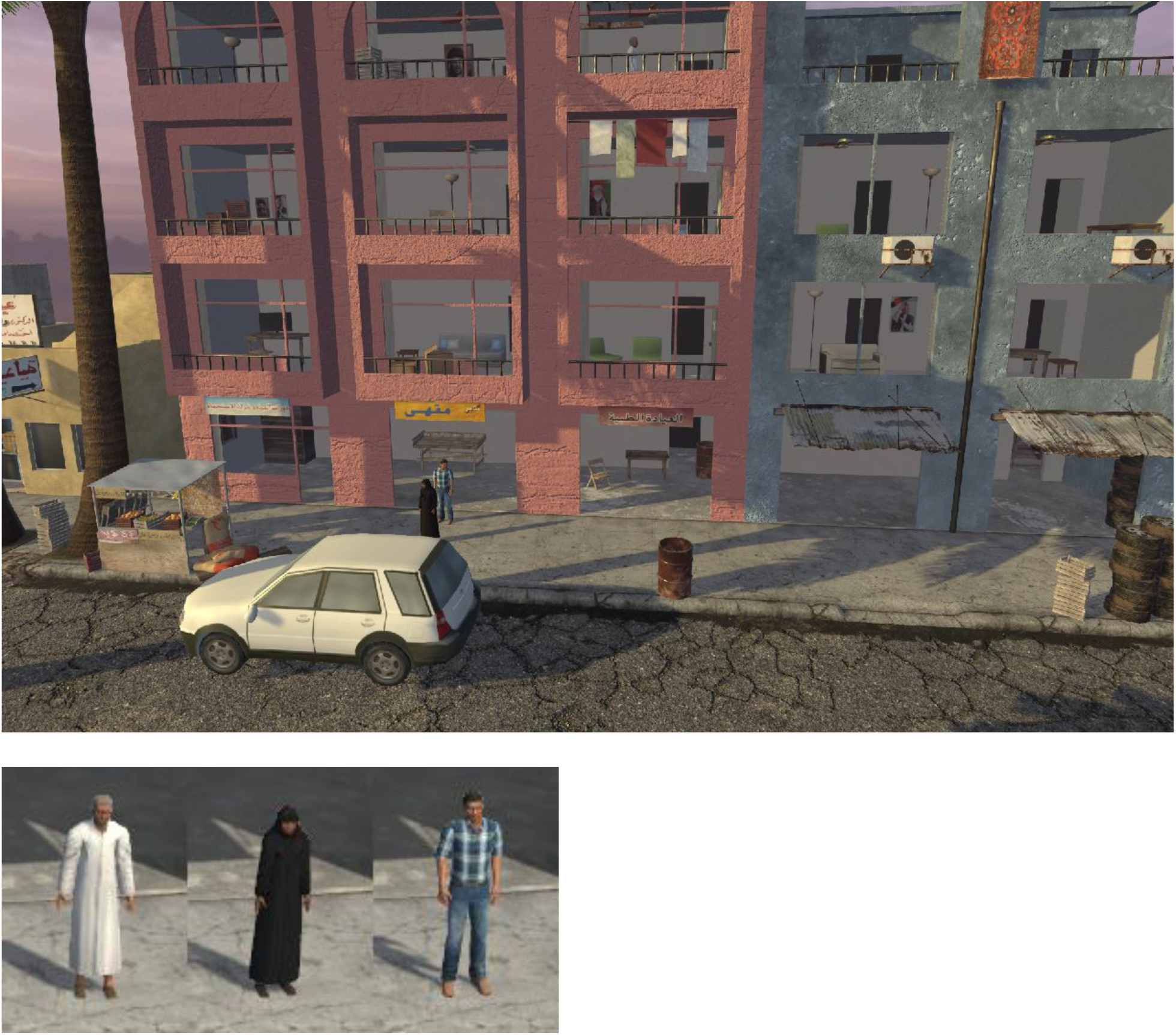

Question 1: Character B was never seen in an upstairs floor prior to the arrival of the car

Answer: True

Question 2: The car arrived after people were already waiting on the sidewalk

Answer: True

Question 3: After running down the stairs Character A gets into the vehicle

Answer: False

Question 4: Character A went back into the building

Answer: False

Question 5: Character A entered the car first

Answer: False

Question 6: Characters B and C exited the building from different exits

Answer: False

Question 7: Characters B and C argued with the driver and the car drove away leaving them on the sidewalk

Answer: False

Question 8: Characters B and C were waiting on the sidewalk when the car drove up

Answer: True

Question 9: There was no indication that Characters B and C knew each other

Answer: True

Question 10: Character A ran down the sidewalk after the car

Answer: True

(16) Missile Strike

Textual prompt: In the episode where two characters are in different rooms of the building when an explosion occurs

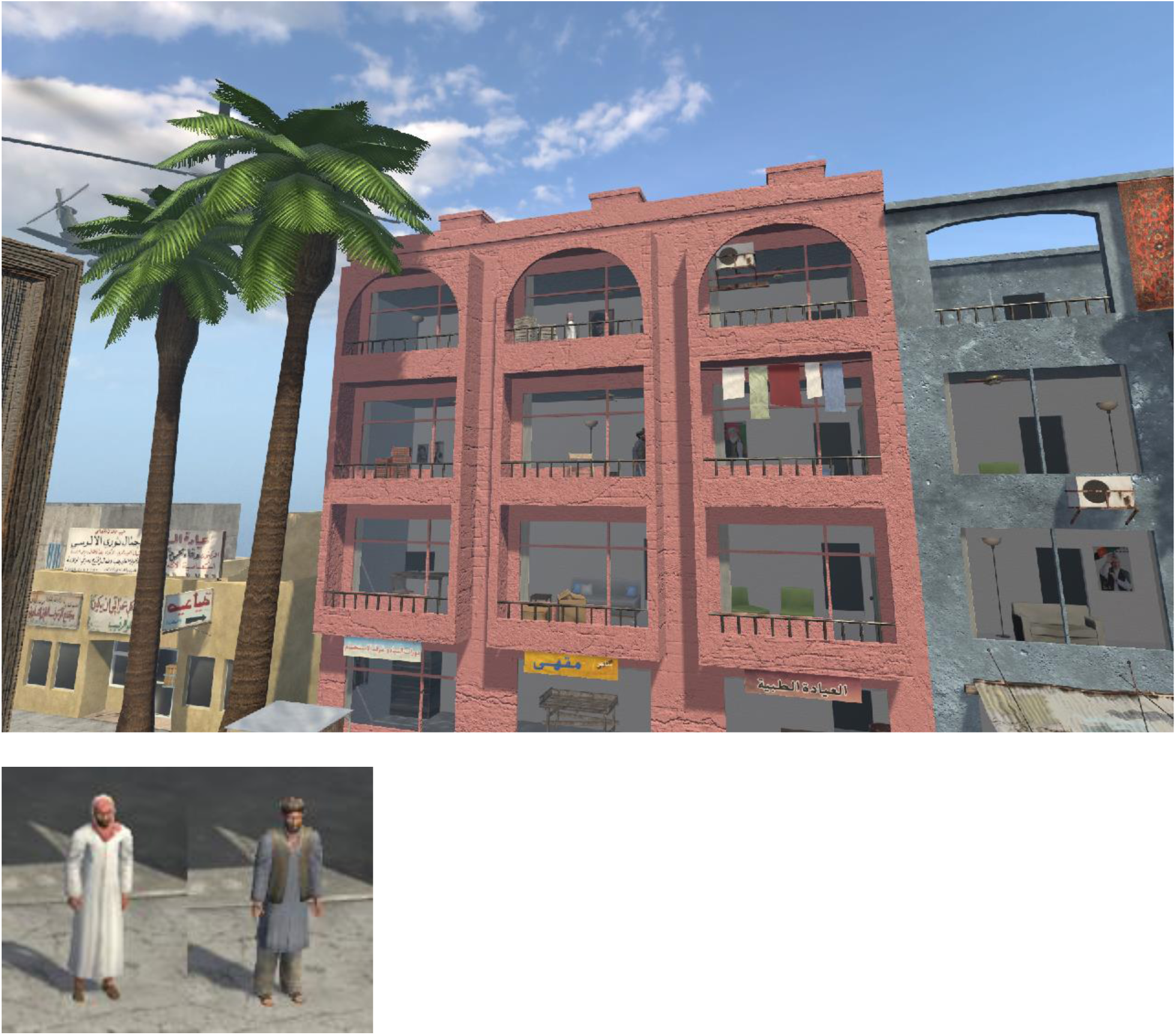

Question 1: Character A enters the building before anyone else was seen in it

Answer: True

Question 2: Character A peeked out of a window before the explosion in that room

Answer: True

Question 3: Character B is visible in the room with the explosion when it occurs

Answer: False

Question 4: Character B ran out of the building right after the explosion

Answer: True

Question 5: An ambulance drives up after explosion

Answer: False

Question 6: A helicopter is heard right before the explosion occurs

Answer: True

Question 7: Character B enters the building after the explosion

Answer: False

Question 8: Character A falls down while running on the street

Answer: False

Question 9: Character A is not seen after explosion

Answer: True

Question 10: Character A fires multiple gunshots before helicopter is heard

Answer: False

(17) No Escape

Textual prompt: In the episode where two characters are in the building when a fire starts and spreads throughout the structure

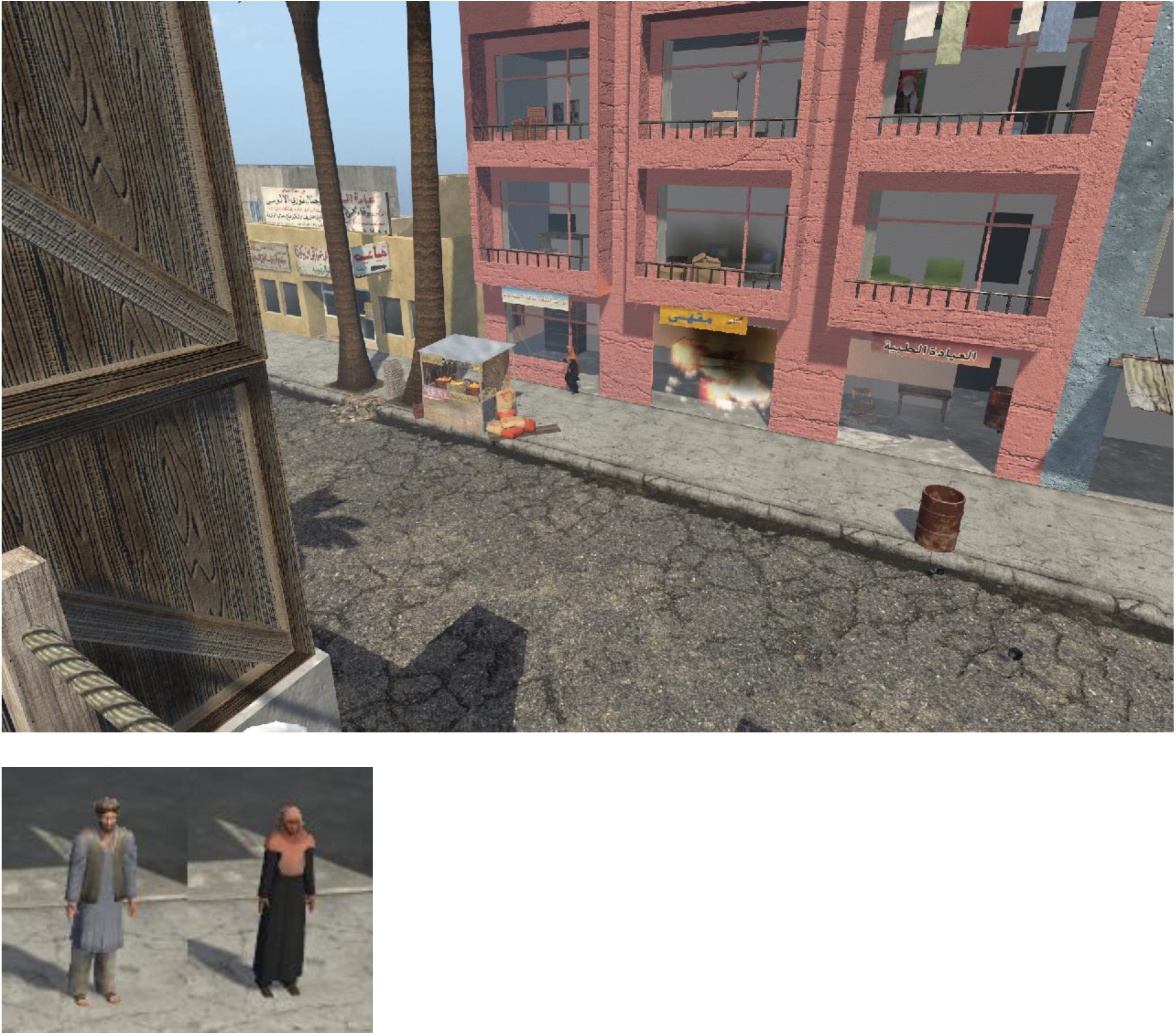

Question 1: An explosion occurs on the bottom floor at the start of the episode

Answer: True

Question 2: Character A runs out of the building after the explosion

Answer: False

Question 3: As the fire spreads, an ambulance arrives

Answer: False

Question 4: The fire spreads across the bottom floor before spreading upward

Answer: True

Question 5: Fire eventually spread to all floors of the building

Answer: True

Question 6: Character B is first seen the 3rd floor, and later is seen on the top floor

Answer: False

Question 7: Character A falls down in a room that is on fire

Answer: True

Question 8: Character B reenters the building after leaving it

Answer: False

Question 9: Characters A and B meet in the building before the fire

Answer: False

Question 10: Character A is seen on the top floor

Answer: True

(18) Pick Pocket

Textual prompt: In the episode where one character takes something from the other while their back is turned, and then a chase ensues down the sidewalk

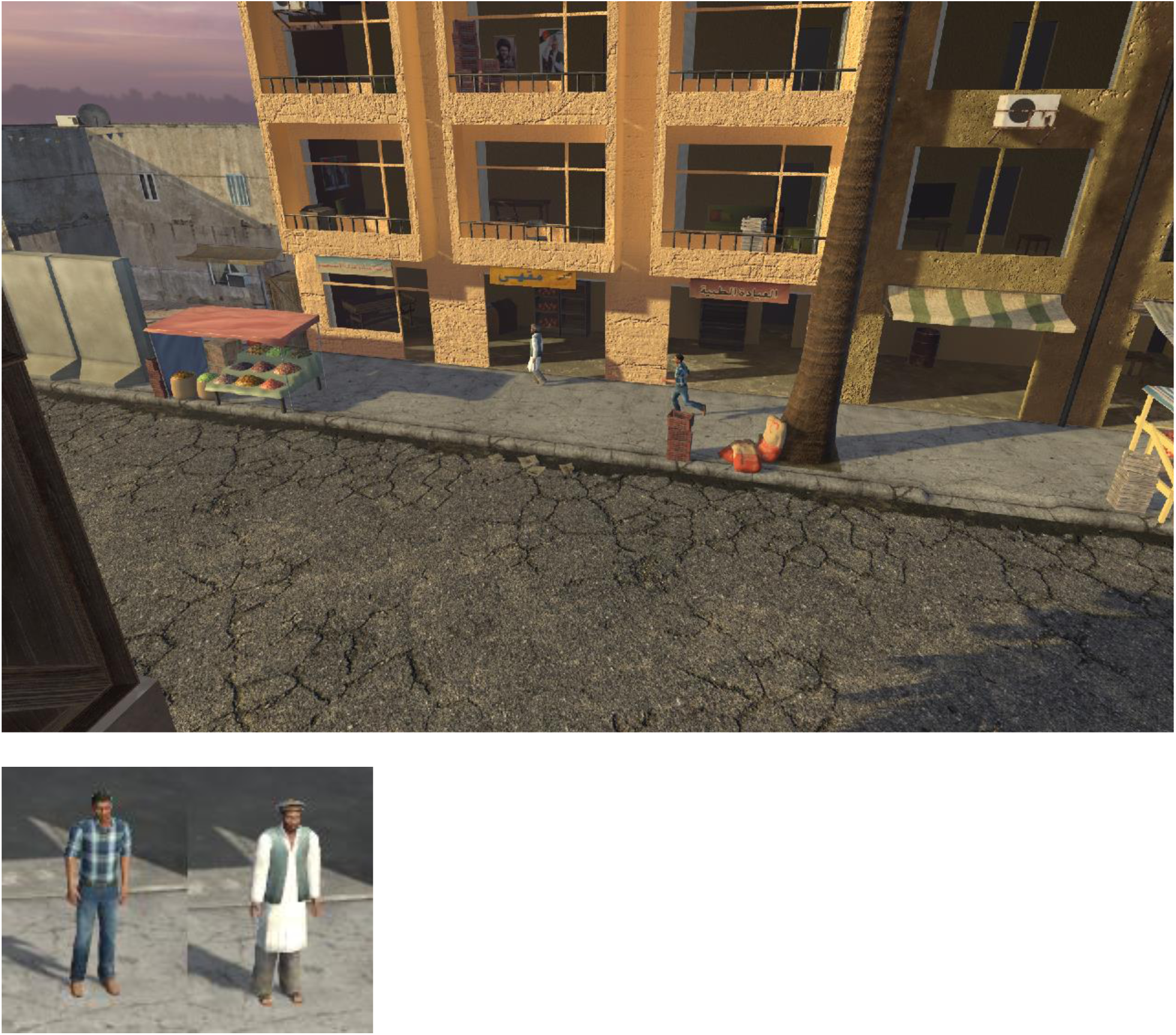

Question 1: Character B faces the building after leaving it

Answer: True

Question 2: Character A got out of a car and stood close to Character B

Answer: False

Question 3: Characters A and B continue running in the same direction

Answer: True

Question 4: Character B entered the building

Answer: False

Question 5: A police car arrived

Answer: False

Question 6: Character B shoots at Character A

Answer: False

Question 7: Character B is not seen catching Character A

Answer: True

Question 8: Character A followed Character B down the sidewalk

Answer: True

Question 9: Character A jumped in a car and drove off

Answer: False

Question 10: Character B walks by Character A before Character A takes something from Character B

Answer: True

(19) Repair Man

Textual prompt: In the episode where smoke is coming from a car by the sidewalk until a character fixes it

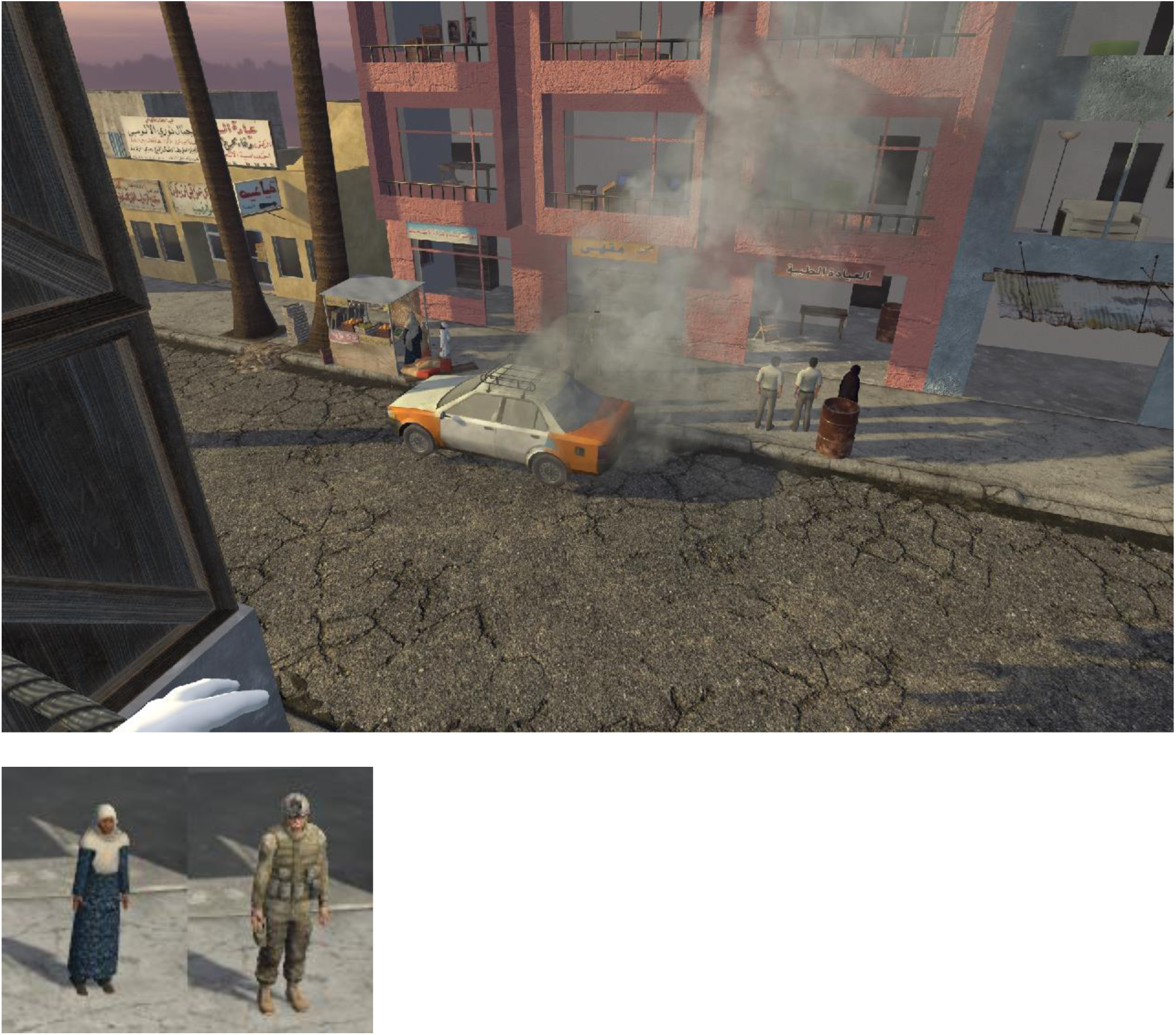

Question 1: The car is already parked by the sidewalk at the start of the episode

Answer: False

Question 2: Character B approaches the car from the left

Answer: True

Question 3: Characters A and B both enter the car

Answer: False

Question 4: Character A goes into the building while Character B is interacting with the car

Answer: False

Question 5: Character A drives off in the car after it stops smoking

Answer: True

Question 6: Character A gets out the car after it starts smoking

Answer: True

Question 7: Character B goes into the building with Character A

Answer: False

Question 8: Character B joins Character A on the sidewalk while the car is smoking

Answer: True

Question 9: Character B steps toward the car and appears to take something out of his pocket, and then the smoke disappears

Answer: True

Question 10: Characters A and B converse again after the car stops smoking

Answer: True

(20) Shopping

Textual prompt: In the episode where several characters enter the shop, some speak to the owner, and some leave

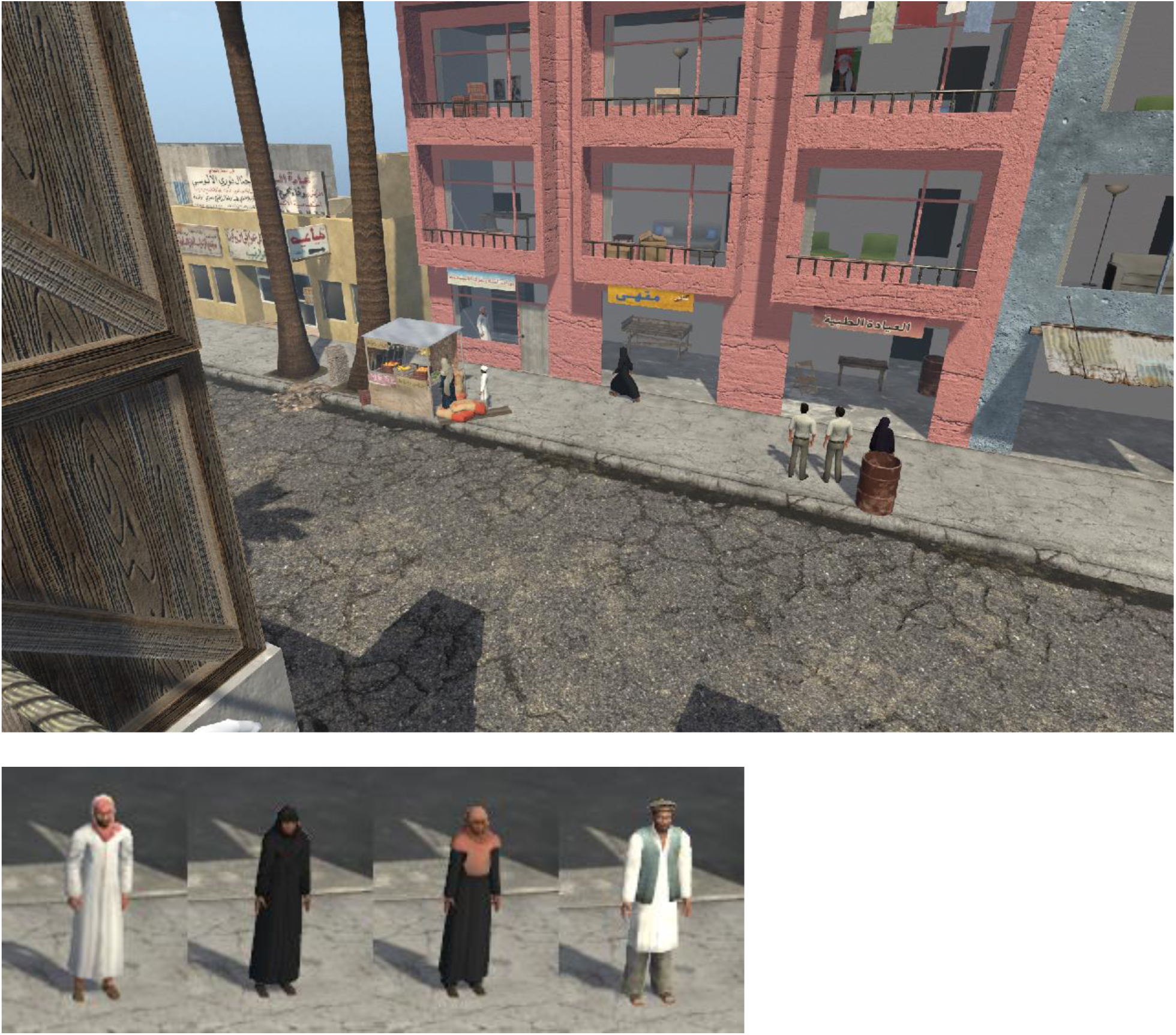

Question 1: Character B got out of a truck and went into a shop

Answer: False

Question 2: Character B paused to look in a window before entering the shop

Answer: True

Question 3: Character A shot Character B in the shop

Answer: False

Question 4: Character C entered the room Character B had left

Answer: True

Question 5: Character D left the building and drove away in a car

Answer: False

Question 6: Character C got out of a car and entered the shop

Answer: True

Question 7: Characters C and D got out of the same car

Answer: False

Question 8: Character D fell down on the sidewalk outside

Answer: False

Question 9: Character A remains in the shop as Character C enters

Answer: True

Question 10: Character B is seen speaking with the owner

Answer: True

(21) Sniper

Textual prompt: In the episode where two characters are conversing on the sidewalk while another is on the roof and gunshots are heard

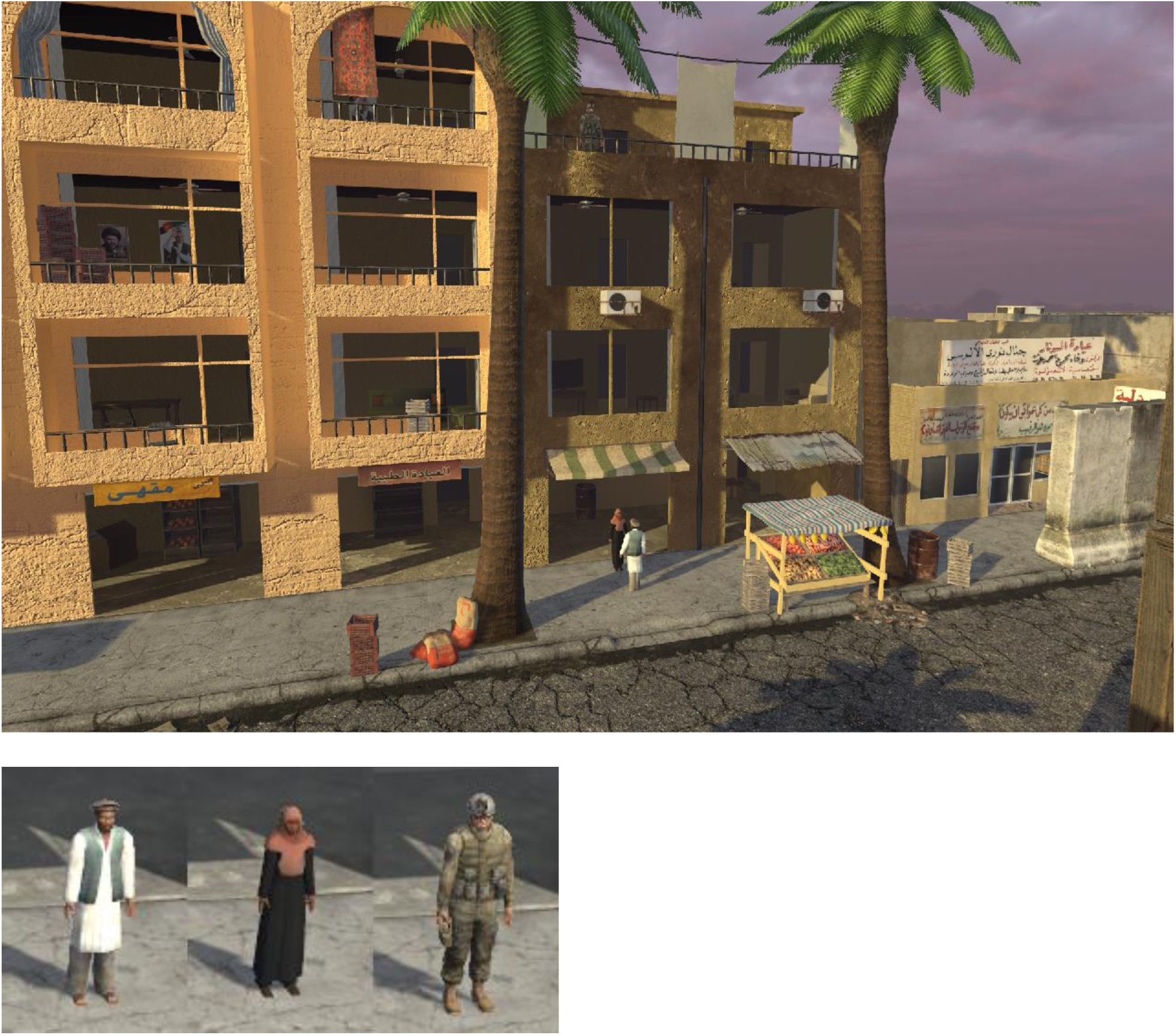

Question 1: Character B ran away alone after gunshots were heard

Answer: True

Question 2: Character B ran into the building after shots were heard

Answer: False

Question 3: Characters A and B ran away when Character C appeared on roof

Answer: False

Question 4: Character C appeared on the roof, and then disappeared and automatic gunfire was heard

Answer: True

Question 5: After the shots were heard, Character C reappeared on the roof

Answer: True

Question 6: Character A got shot and Character B ran down the sidewalk to the left

Answer: True

Question 7: Character B left more quickly than they arrived

Answer: True

Question 8: An ambulance arrived

Answer: False

Question 9: Characters A and B came out of the building

Answer: False

Question 10: Character B got in a car and drove away

Answer: False

(22) Street Hawker

Textual prompt: In the episode where one character on the sidewalk stops the other to talk, follows them, and then convinces them to enter the building

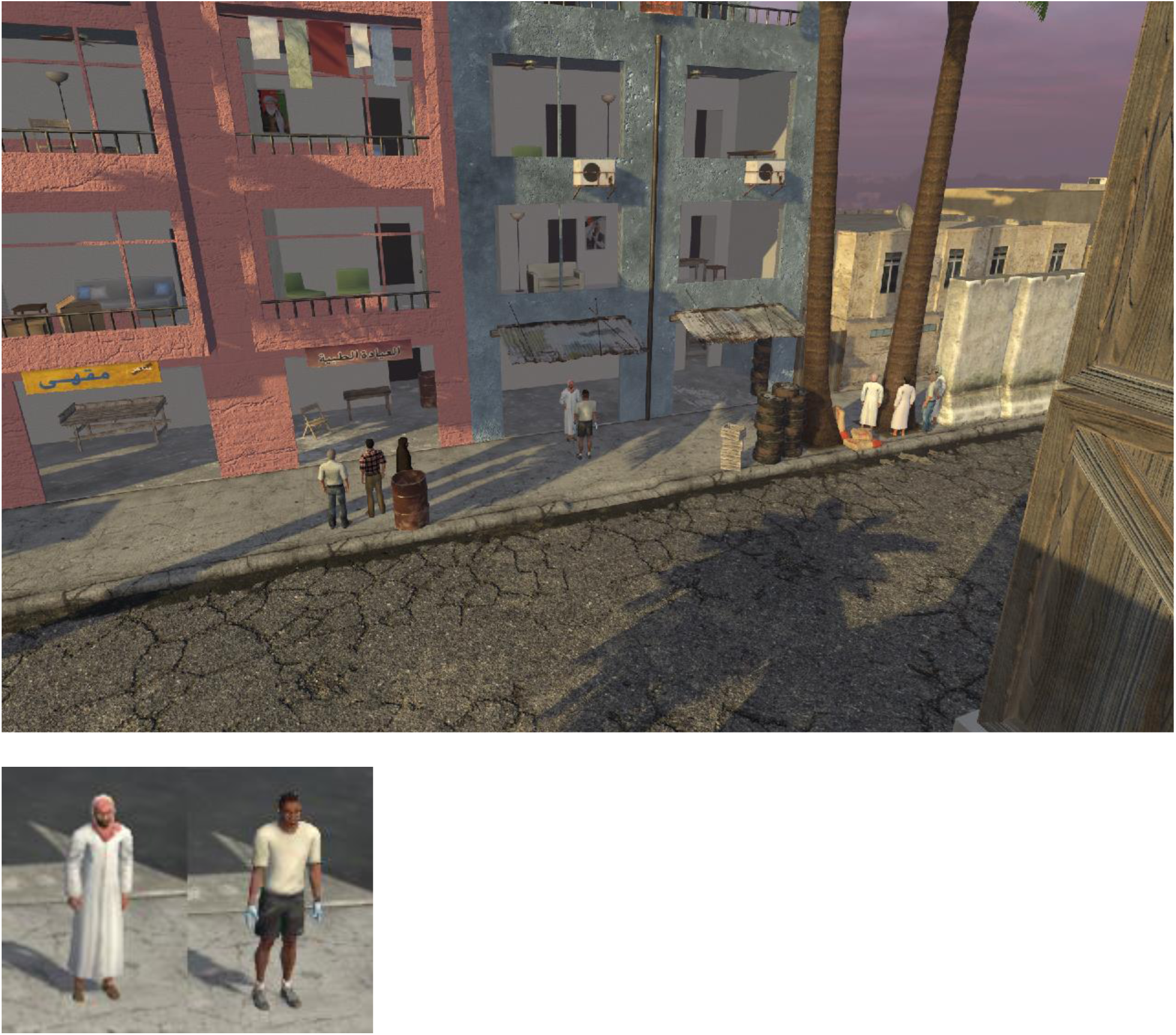

Question 1: Two people met on the sidewalk at right side of building, then walked to left side of building

Answer: True

Question 2: After stopping at left side of building, Character A got in a car and drove away

Answer: False

Question 3: Characters A and B conversed a second time after Character B walks further down the street

Answer: True

Question 4: Gunshots are heard before the two enter the building

Answer: False

Question 5: The people talking on the sidewalk entered the building on the right side, and the garage door closed behind them

Answer: True

Question 6: Smoke is seen coming from the room the characters entered

Answer: True

Question 7: An emergency vehicle arrives after smoke is seen

Answer: False

Question 8: Characters A and B enter the building together

Answer: True

Question 9: Characters A and B converse in a room on an upper floor of the building

Answer: False

Question 10: Character A enters the building while Character B remains outside

Answer: False

(23) Suicide Bomb

Textual prompt: In the episode where one character enters the building, meets with the other character in a room, then there is an explosion in that room

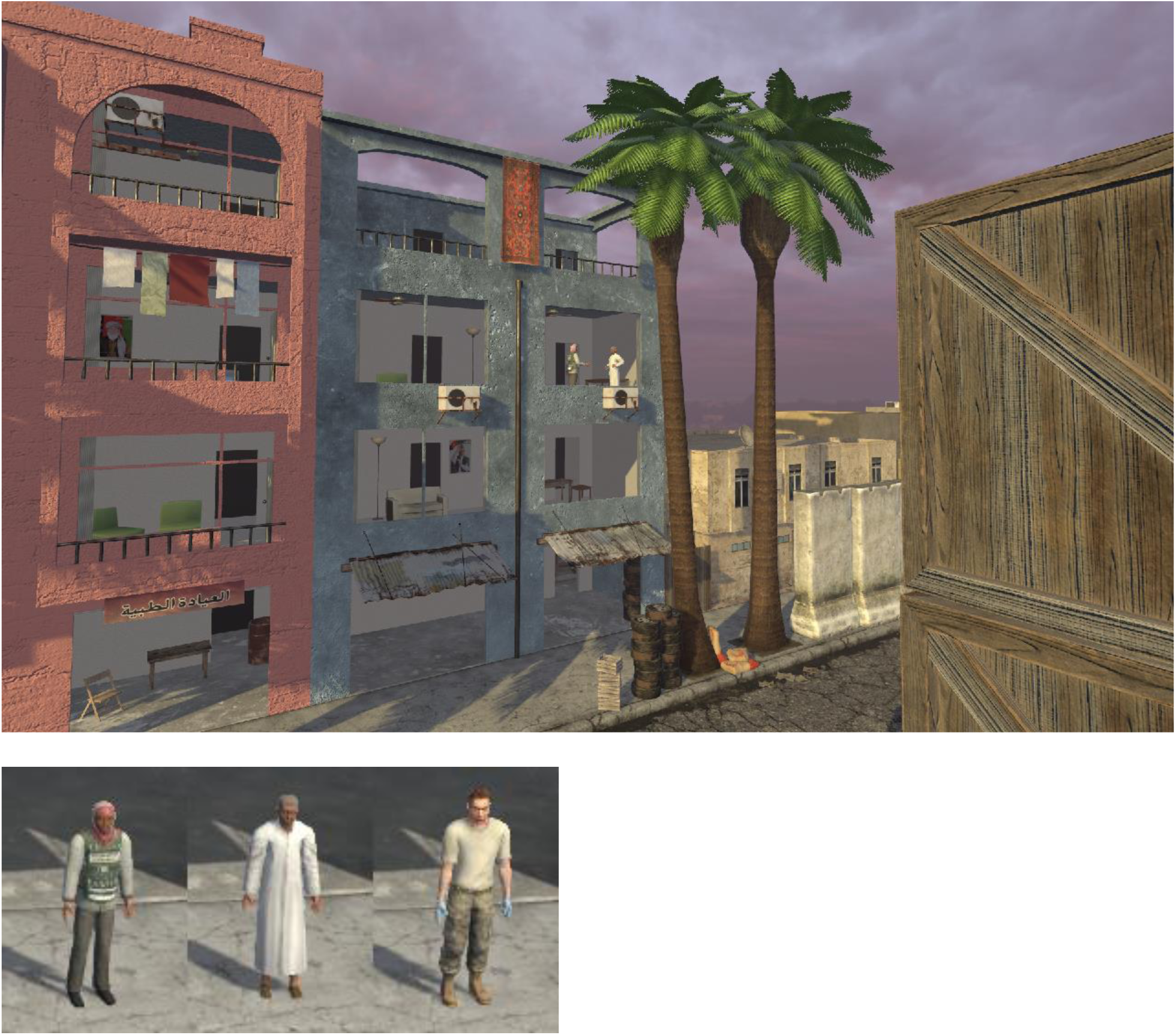

Question 1: Character A entered the left entrance of the building

Answer: False

Question 2: Character B was already in the building when Character A entered

Answer: True

Question 3: An ambulance arrives after explosion

Answer: True

Question 4: Character B tries to run before the explosion occurs

Answer: False

Question 5: Fire starts in the room after the explosion

Answer: True

Question 6: Character A runs from building after the explosion

Answer: False

Question 7: Character C arrives in a vehicle

Answer: True

Question 8: Character B leaves in a car

Answer: False

Question 9: Characters A and B are not seen after the explosion

Answer: True

Question 10: Characters A and B converse outside after the explosion

Answer: False

(24) The Dare

Textual prompt: In the episode where two characters start a fire in the building

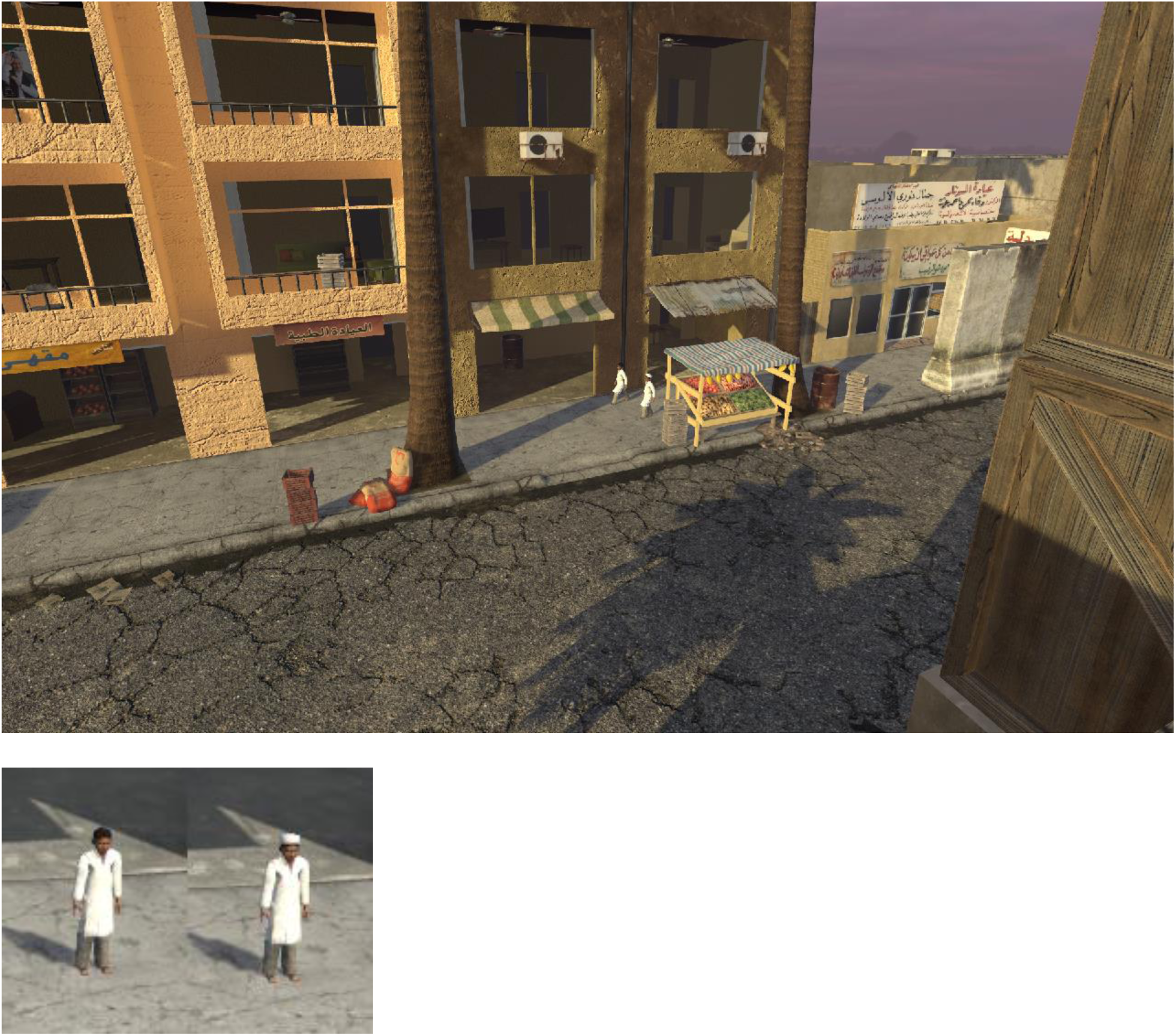

Question 1: Characters A and B stop to talk outside building

Answer: True

Question 2: Characters A and B enter the building together

Answer: False

Question 3: After Character B enters the building a room catches fire

Answer: True

Question 4: A police vehicle drives in the direction Characters A and B ran away

Answer: False

Question 5: Character B leaves the building after the fire starts

Answer: True

Question 6: Character B passes through a room while it is on fire

Answer: False

Question 7: Character A waits outside while Character B enters the building

Answer: True

Question 8: Character B gets in a car and drives off

Answer: False

Question 9: The fire spreads to many rooms

Answer: False

Question 10: Characters A and B leave headed the same direction

Answer: True

(25) The Meeting

Textual prompt: In the episode where two characters meet in a room and a third character is seen on the sidewalk

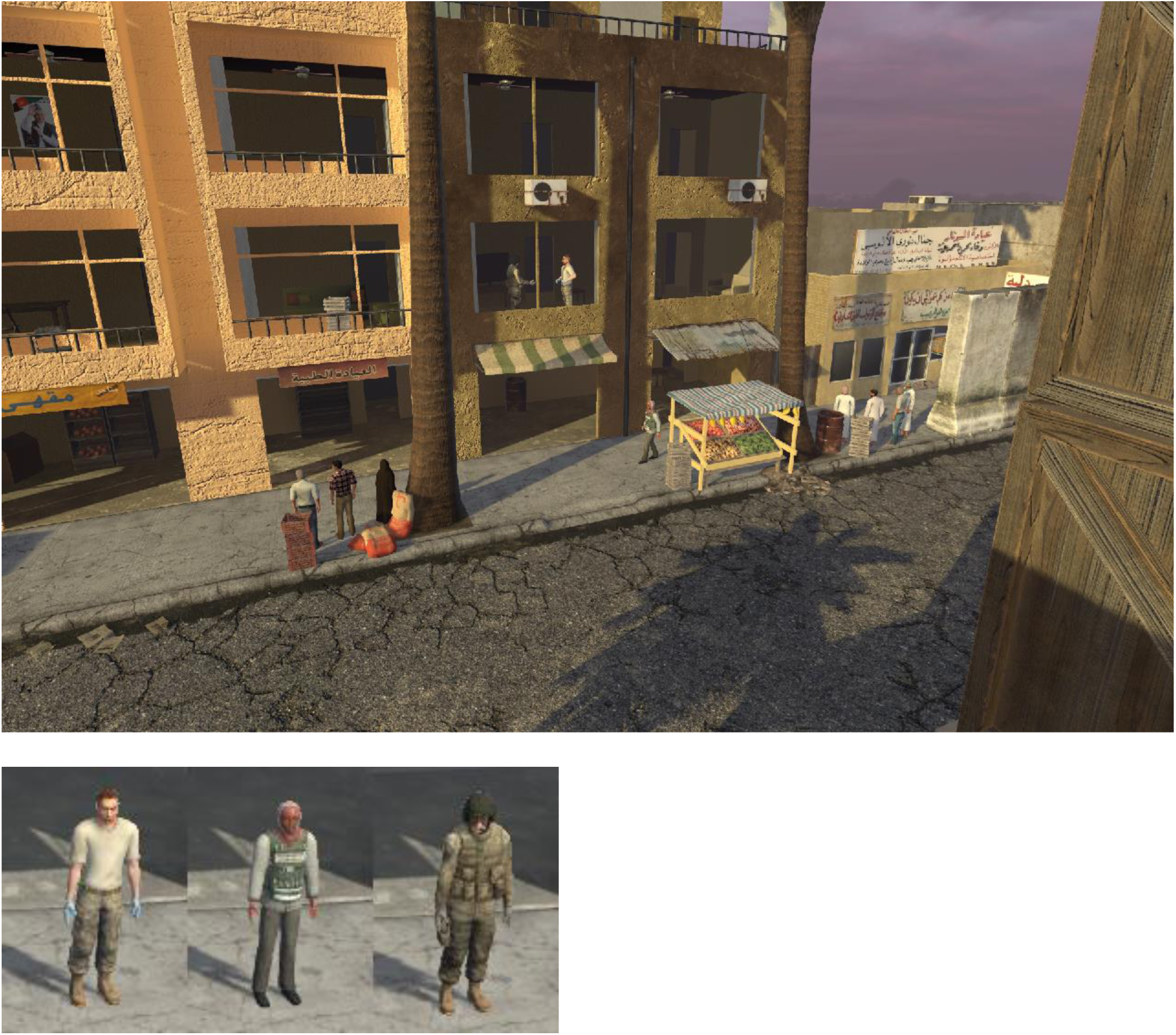

Question 1: Character A enters the building before Character C

Answer: True

Question 2: Character B came out of the building

Answer: False

Question 3: Characters A and C leave the building and go the same direction as Character B

Answer: True

Question 4: Character B enters the building through the same entrance as Characters A and C

Answer: False

Question 5: Character A watches out the window as Character C approaches

Answer: True

Question 6: Characters A and C close the curtains before they leave the room

Answer: False

Question 7: Character B got out of a car

Answer: False

Question 8: A police car drove by

Answer: False

Question 9: Characters A and C both entered the same building entrance

Answer: True

Question 10: Character B ran away when he saw the other two in the building

Answer: True

(26) The Serenade

Textual prompt: In the episode where one character on the sidewalk sings to the other character in the building

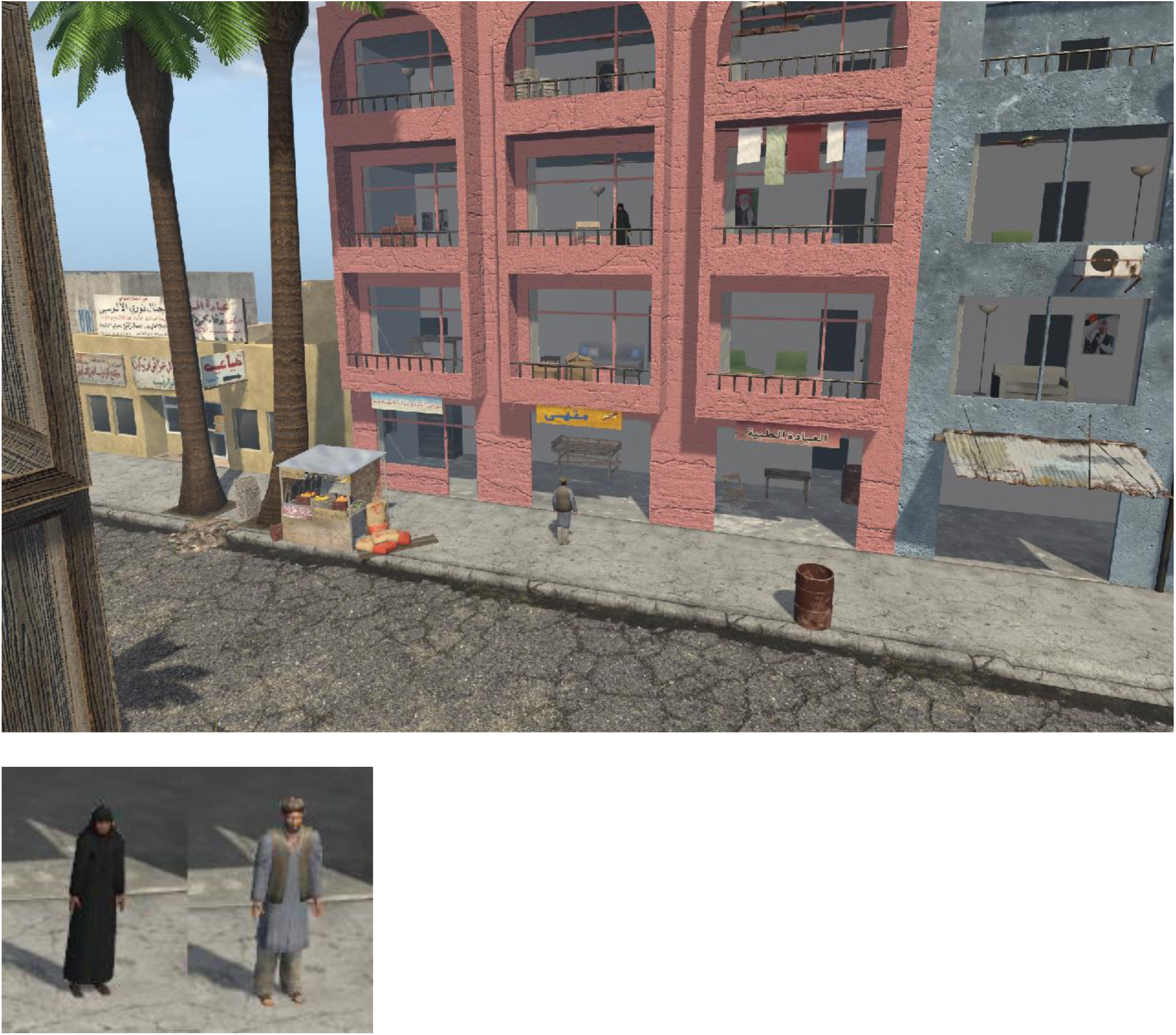

Question 1: Character B came walking down the sidewalk from the right to the front of the building

Answer: True

Question 2: Character A waited for Character B to finish singing before coming down

Answer: True

Question 3: Character A in the building drew the blinds after listening

Answer: False

Question 4: Character B started singing before Character A came to the window

Answer: True

Question 5: Character A came down and joined Character B, and they went down the sidewalk together

Answer: True

Question 6: Character B arrived in a car

Answer: False

Question 7: Character A came down and joined Character B, and they left in a car

Answer: False

Question 8: Character B entered the building and met Character A in her room

Answer: False

Question 9: There was no vehicle of any kind in this vignette

Answer: True

Question 10: After speaking on the side walk the two characters walked in opposite directions

Answer: False

(27) Timely Exit

Textual prompt: In the episode where one character speaks with the other character in the building, and they leave before an explosion occurs in the room they were just in

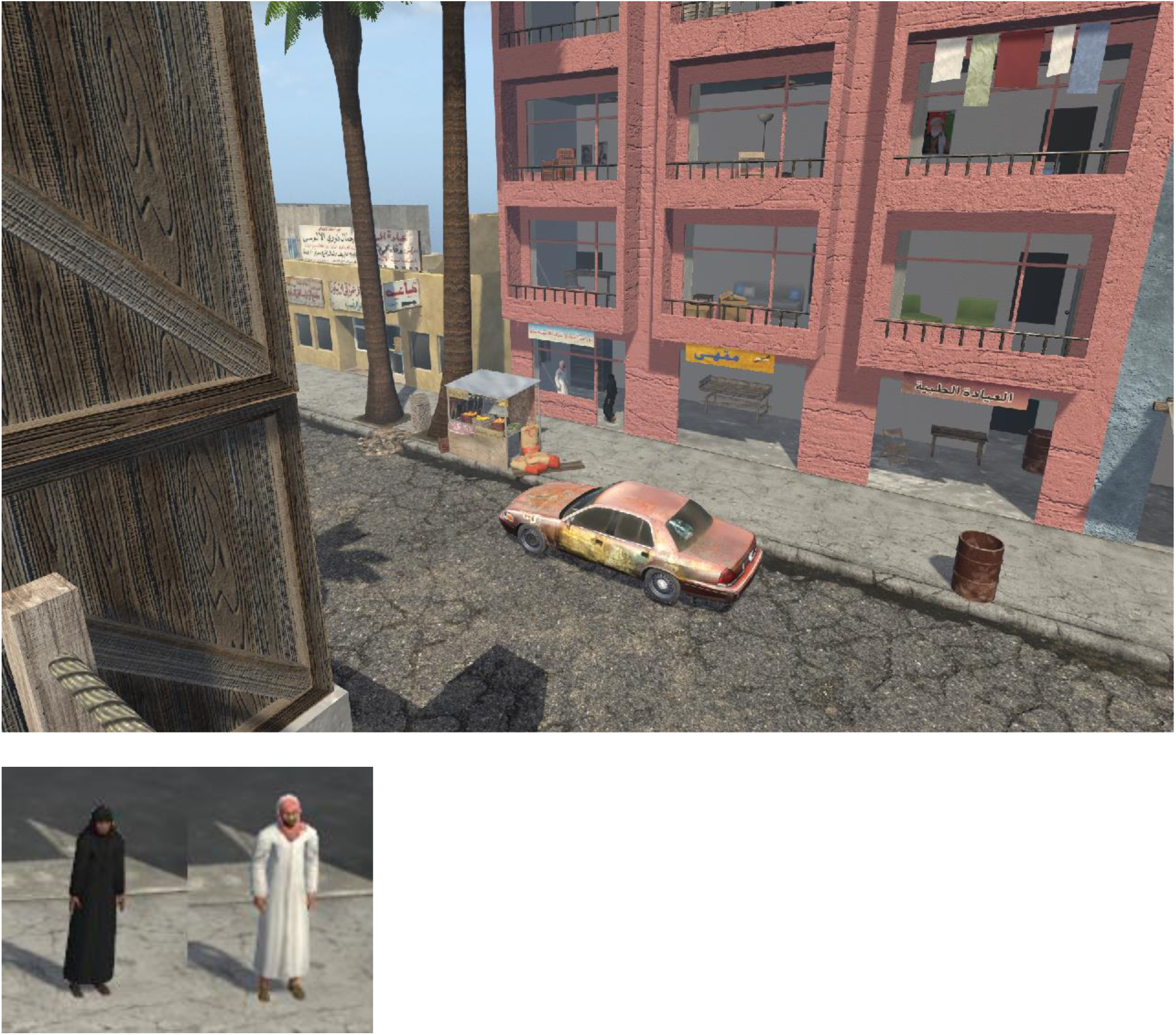

Question 1: Character A arrived in a car

Answer: True

Question 2: Character A entered the building to converse with Character B

Answer: True

Question 3: The door of the store was open while the two people talked inside

Answer: False

Question 4: A second vehicle arrives before first leaves

Answer: False

Question 5: There was a second explosion after the first

Answer: False

Question 6: A helicopter flew over just before the explosion

Answer: True

Question 7: An ambulance arrives after the explosion

Answer: False

Question 8: Character A falls down in the street after explosion

Answer: False

Question 9: Character B runs down the street after explosion

Answer: False

Question 10: Characters A and B entered a car and drove away

Answer: True

(28) Waiting Gunman

Textual prompt: In the episode where one character shoots the other as they leave the building

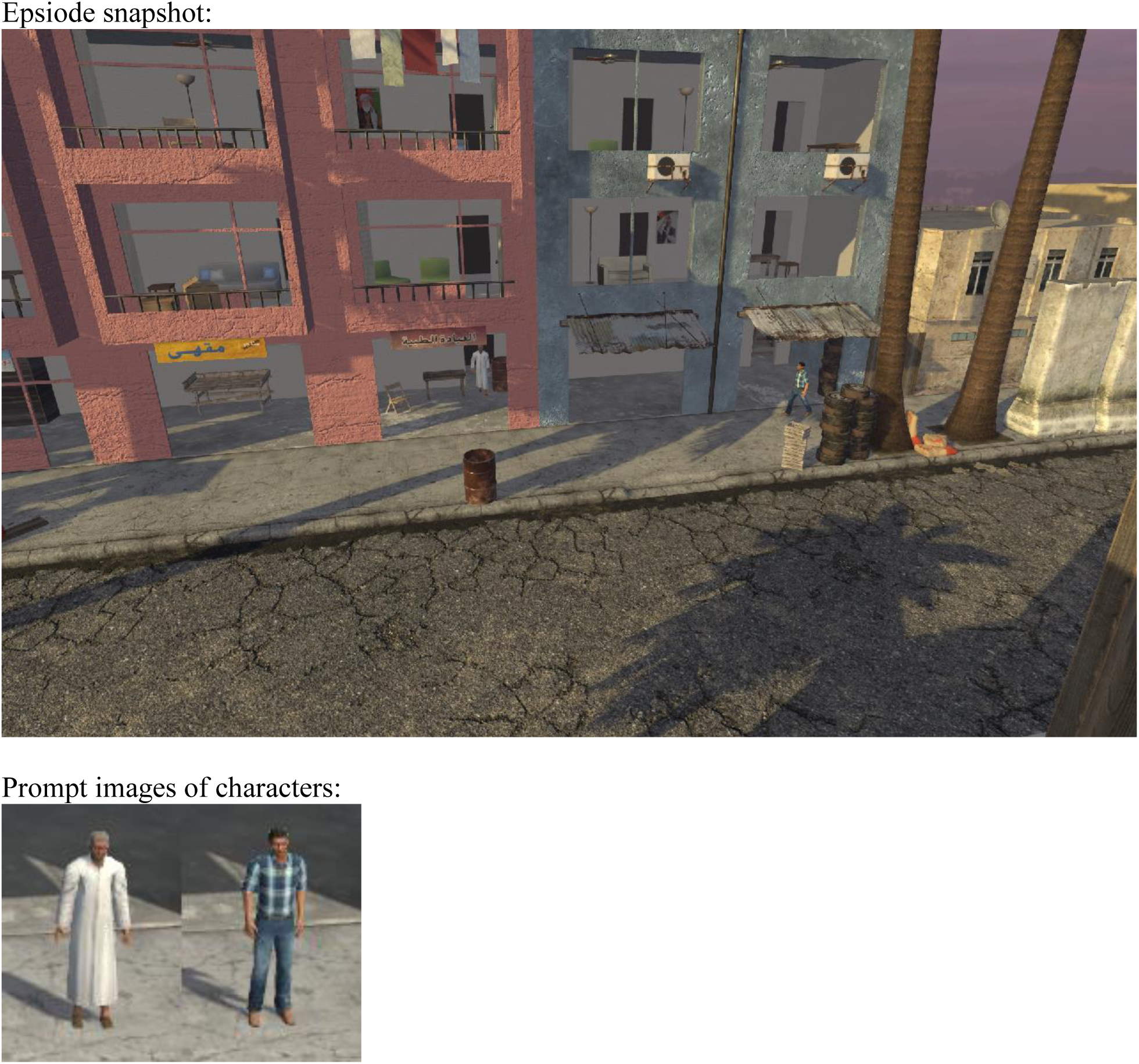

<u>The 10 questions:</u>

Question 1: Character B fires many shots

Answer: True

Question 2: Character B shoots at Character A

Answer: True

Question 3: Character A falls while running away

Answer: True

Question 4: Character B runs past Character A

Answer: True

Question 5: A car pulls up and Character B gets in

Answer: False

Question 6: Character A starts to run when shooting begins

Answer: True

Question 7: Characters A and B are facing each other when shots are fired

Answer: False

Question 8: Character A looks out of a window before leaving the building

Answer: False

Question 9: An emergency vehicle arrives

Answer: False

Question 10: Character B turns to run away from the place where Character A has fallen

Answer: False

